# Accurate Protein Dynamic Conformational Ensembles: Combining AlphaFold, MD and Amide ^15^N(^1^H) NMR Relaxation

**DOI:** 10.1101/2025.02.07.637034

**Authors:** Dmitry Lesovoy, Konstantin Roshchin, Benedetta Maria Sala, Tatyana Sandalova, Adnane Achour, Tatiana Agback, Vladislav Orekhov, Peter Agback

## Abstract

Conformational heterogeneity is critical for protein function, but the validation of dynamic ensembles remains a challenge. In this study, we introduced an approach that integrates free MD simulations, using an AlphaFold-generated structure as the starting point, with experimental relaxation data to identify biologically relevant conformational ensembles. For the extracellular region of *Streptococcus pneumoniae* Psr_Sp_, we found that only certain segments of the MD long trajectory aligned well with experimental data. The defined ensembles revealed two regions with increased flexibility that play important functional roles.

## Introduction

Over the past decade, conformational ensembles have gained increasing recognition as the most accurate representation of a protein’s native state, offering valuable insights into the fundamental relationships between protein structure, dynamics, and function (Friedland *et al*., 2009, Boehr *et al*., 2009). This shift came from by the realization that traditional paradigms fail to fully capture the complexity of biological functions, as they neglect the dynamic nature of proteins (Nussinov, 2016, Nussinov, Liu, *et al.*, 2023b). Recent advances in physics and chemistry, particularly the development of energy landscape theory, have significantly reshaped molecular biology by highlighting that proteins continuously fluctuate between multiple conformational states, each corresponding to distinct energy levels (Nussinov, Liu, *et al.*, 2023b). This dynamic perspective offers a more realistic and comprehensive understanding of protein function in living systems.

For decades, the relationship between sequence, structure, and function in molecular biology was based on the assumption that each protein sequence folds into a single, averaged structure under given conditions. This foundational belief deeply influenced traditional structural biology approaches and is reflected in widely used software packages for nuclear magnetic resonance (NMR) spectroscopy, such as CNS (Brunger *et al*., 1998), XPLOR (Schwieters *et al*., 2003), CYANA (Guntert, 2004, Klukowski *et al*., 2024), HADDOCK (Dominguez *et al*., 2003) and CS-RosettaCM (Shen & Bax, 2015), originally designed to produce a single, structure that satisfies all conformationally averaged experimental constraints. Despite its limitations, the single-structure paradigm facilitated the creation of extensive public databases of experimentally determined protein structures, primarily obtained through X-ray crystallography, cryo-electron microscopy (cryo-EM) and NMR spectroscopy (Klukowski *et al*., 2024). Capitalising on this data, groundbreaking developments in Artificial Intelligence, such as AlphaFold, have significantly improved the predictions of static protein structures (Jumper et al., 2021).As structural biology shifts from studying well-defined macromolecules toward larger, more complex, and flexible molecular systems, there is an increasing need for structural approaches capable of capturing the full spectrum of conformational heterogeneity. This transition from <u>static</u>, single-structure models to dynamic ensemble representations requires the development of novel, conceptually distinct computational methods and experimental tools (Shapiro, 2013, Torchia, 2015, van den Bedem & Fraser, 2015, Wei *et al*., 2016, Ravera *et al*., 2016, Dokholyan, 2020, Costa & Fushman, 2022, Ramelot *et al*., 2023, Schwalbe *et al*., 2024).

Solution-state NMR spectroscopy is a powerful tool for studying conformational ensembles as it inherently captures the physical properties of biomolecules averaged across multiple conformations, offering insights into protein dynamics across a wide range of timescales. From the beginning, NMR datasets - such as chemical shifts (CS) (Jensen *et al*., 2010, Robustelli *et al*., 2013, Palmer, 2016), residual dipolar couplings (RDCs) (Clore & Schwieters, 2004, Lindorff-Larsen *et al*., 2005, Markwick *et al*., 2007, Showalter & Bruschweiler, 2007, Lange *et al*., 2008, Nodet *et al*., 2009, Torchia, 2015, Ravera *et al*., 2016, Shen & Bax, 2023), paramagnetic relaxation enhancements (PREs) (Allison *et al*., 2009, Bertini *et al*., 2017) -have been the primary choice for defining conformational ensembles.

Despite NMR’s exceptional ability to probe backbone and side-chain dynamics, relaxation measurements have been relatively underutilized in the determination of structural ensembles. Relaxation measurements, such as longitudinal (R₁), transverse (R₂), and heteronuclear NOE, provide detailed insights into the dynamics structural ensembles, reflecting their heterogeneity and temporal properties (Palmer, 2004, Cavanagh *et al*., 2007, Trbovic *et al*., 2008, Palmer, 2015, Torchia, 2015, Palmer, 2016). Early studies employed the model-free (MF) approach (Kay *et al*., 1989, Korzhnev *et al*., 2001, Zumpfe & Smith, 2021) to estimate the rates and amplitudes of internal motions on the pico- to nanosecond timescale. The analysis yields the generalized order parameter (S²), which quantifies the structural range of fast internal motions, (from 0, indicating complete disorder to 1, indicating complete rigidity), and the correlation time (τ_c_), which reflects the timescale of structural fluctuations. Interpretation of NMR relaxation data in the context of conformational ensembles remains challenging due to the difficulty in distinguishing structural features from dynamic behaviour (Kauffmann *et al*., 2021, Stenstroem *et al*., 2022). However, recent advances in molecular dynamics (MD) simulations-driven by improved force fields (Dauber-Osguthorpe & Hagler, 2019, Schlick & Portillo-Ledesma, 2021, Schlick *et al*., 2021, Zhang *et al*., 2022, Kang *et al*., 2022) and more affordable access to high-performance computing-have enabled the integration of relaxation data with computational models. This integration allows for more accurate modelling and validation of dynamic conformational ensembles sampled on the picosecond-to-nanosecond timescale. Several strategies have been developed to combine NMR relaxation data with MD simulations to capture dynamic conformational states in solution. The original approach employs constrained MD simulations with additional force-field terms to obtain MD trajectories aligned with the experimental model-free order parameters and other NMR data (Lindorff-Larsen *et al*., 2005, Torchia, 2015, Orioli *et al*., 2020).

Another method extracts backbone **¹H–¹⁵N** vector motions from an unconstrained MD trajectory calculated with the most realistic force fields, followed by back-calculation of order parameters or NMR relaxation rates (Pfeiffer *et al*., 2001, Nederveen & Bonvin, 2005, Showalter & Bruschweiler, 2007, Salvi *et al*., 2017, Kämpf *et al*., 2018, Salvi *et al*., 2022, Agback *et al*., 2023). Various MD force fields were benchmark using-model proteins like ubiquitin against the experimental **R₁**, **R₂**, and **NOE**-derived order parameters (**S²**) (Koller *et al*., 2008, Showalter & Bruschweiler, 2007, Kummerer *et al*., 2023).

Along this line, Palmer and colleagues (Robustelli *et al*., 2013) were probably the first to demonstrate that NMR relaxation data can serve to select MD trajectories inconsistent with experimental dynamics. They showed that back-calculated NMR chemical shifts and spin-relaxation data provide complementary insights into the structure and dynamics of intrinsically disordered proteins (IDPs). Their work revealed a strong agreement between experimental and computed generalized order parameters, allowing the identification of MD trajectories that most accurately reflect experimental observations. This approach for exploring conformational ensembles in IDP (Kämpf *et al*., 2018) and global proteins (Agback *et al*., 2023, Banayan *et al*., 2024) has since been applied..

The analysis was further improved by replacing experimental **R₂**, which may be biased by the slow conformational exchange, with the cross-correlated relaxation (H2) rates (Kämpf *et al*., 2018).

Building on this progress, we previously conducted a study validating dynamic ensembles of the Dengue II protease protein derived from unconstrained MD simulations (Agback *et al*., 2023). In that work, we selected MD trajectories by comparing experimental and back-calculated relaxation parameters, including backbone R₁, NOE, R₂, and various types of cross-correlated relaxation in methyl side-chain dynamics. Additionally, we examined how starting molecular models, obtained from experimental methods such as X-ray crystallography and NMR-refined structures of the Dengue II protease, can serve for further refinement, structural analysis, and differentiation between conformational states.

Significant challenges remain in validating theoretical structural-dynamic ensembles, primarily due to incomplete sampling of the conformational space (Orioli *et al*., 2020, Kummerer *et al*., 2023). Recent studies have shown that AlphaFold holds great promise not only in predicting the “best” single structure but also in generating conformational ensembles consistent with experimental and evolutionary data (Kaczmarski *et al*., 2020, Nussinov, Liu, *et al.*, 2023a, Wallerstein *et al*., 2024, Wayment-Steele *et al*., 2024). AlphaFold-generated structural ensembles are considered promising starting points for MD simulations (Heo & Feig, 2020, Ma *et al*., 2023), as they may effectively explore a broad range of local and global energy minima.

Building on these advancements, we present an efficient AlphaFold-MD-NMR based method that uses back-calculated R1, NOE, and H2 relaxation parameter for generation and validation of the dynamical conformation ensembles best aligned with the experimental relaxation data. We also introduce an improved experimental scheme for measuring H2 relaxation in backbone HN groups.

We applied this approach to the extracellular region of *Streptococcus pneumoniae* protein PsrSp (residues 131-424). Psr_Sp_ catalyses the attachment of cell wall teichoic acid and/or other polysaccharides to the peptidoglycan layer of Gram-positive bacterial cell walls, a process critical for bacterial survival (Stefanovíc *et al*., 2021). Recent combined crystallographic and NMR studies of Psr_Sp_ and related LCP homologs highlighted the importance of loops surrounding the active site for substrate binding (Sandalova *et al*., 2024). Variability in loop length and composition across species contributes to substrate specificity, with dynamic conformational changes likely playing a key functional role. Since these regions represent promising targets for antibiotic development, a deeper understanding of their flexibility and dynamics, particularly in the hotspot regions, is essential for guiding future drug discovery efforts. Here, we constructed and validated a structural-dynamic model of Psr_Sp_. Our model reveals the functional mobility of two key hotspot regions: a loop at the active site and a substrate-binding pocket composed of an α-helix and an adjacent irregular segment. The resulting dynamic conformational ensemble depicts how these dynamic regions influence ligand interactions.

Our work signifies a new approach in NMR structural biology, enabling the generation of fully experimentally validated dynamic conformational ensembles using readily accessible NMR relaxation data, without requiring for significantly more costly and time-consuming experimental structure determination methods such as X-ray crystallography, traditional NMR spectroscopy, *etc*.

## Material and Methods

### Sample preparation

All NMR experiments were performed on a U-[^15^N,^13^C,^2^H] labelled sample of Psr_Sp_. The sample preparation has been fully described in our earlier publication (Sandalova *et al*., 2024). In short, the Psr_Sp-_ sequence was cloned into a pET28 vector in frame with a N-terminal His-tag such that 12 amino acid residues (MHHHHHHENLYF) were added; thus, the construct contains 308 residues and have a molecular mass of 35.5kDa (**Figure 1(a)**). The plasmids were transformed into BL21 (DE3) pLys *E. Coli* strain and cells were cultured at 37°C in isotope ^2^H, ^15^N, ^13^C-labelled M9 medium. Chemicals for isotope labelling (ammonium chloride, ^15^N (99%), D-glucose, ^13^C (99%) and deuterium oxide were purchased from Cambridge Isotope Laboratories, Inc.

**Figure 1.**
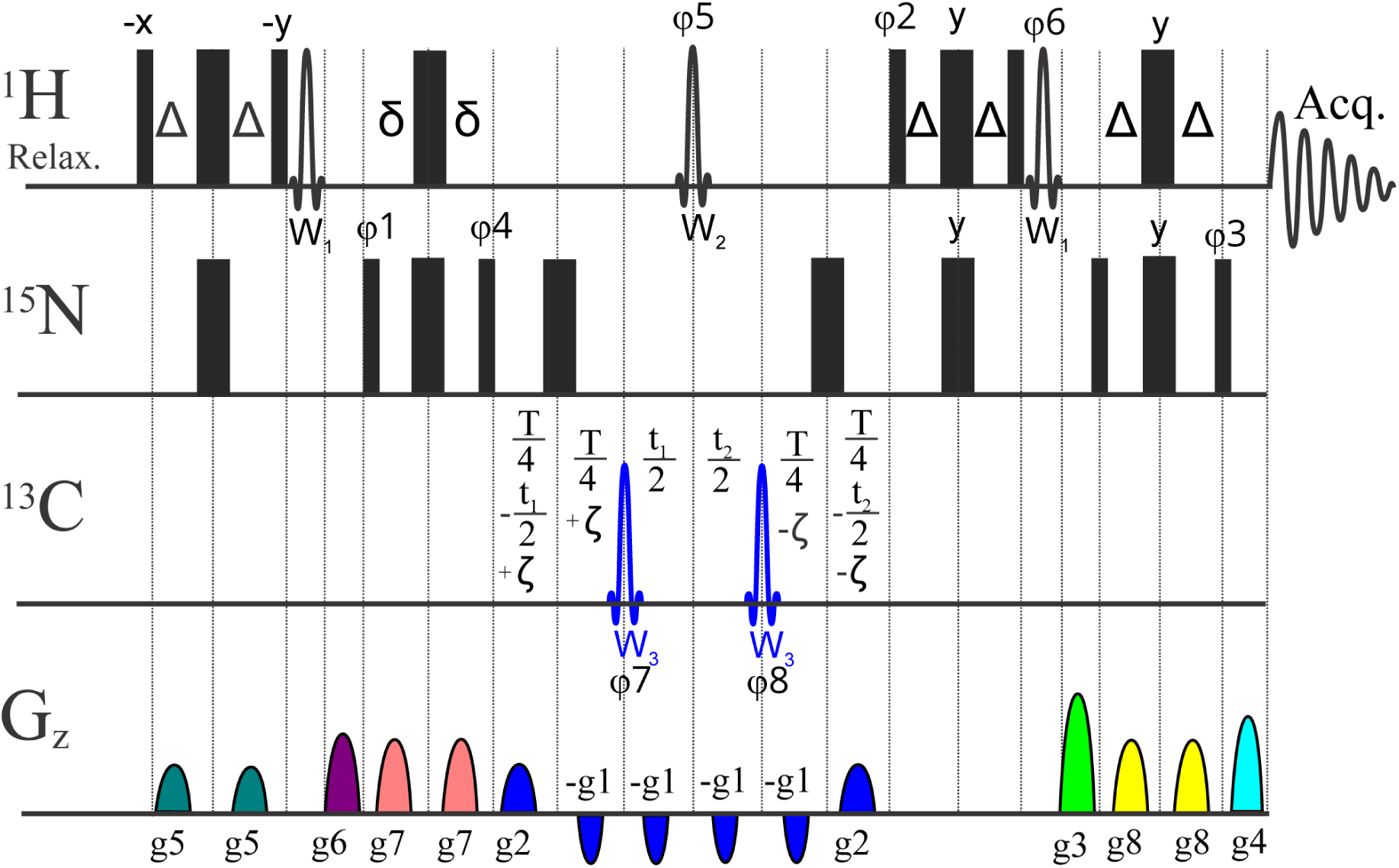
The pulse sequence of pseudo 3D ^1^H-^15^N proton-detected experiment for measurement of ^1^H-^15^N CSA/DD transverse cross correlation relaxation in backbone ^15^N^1^H groups. Narrow and thick bars represent 90° and 180° pulses respectively, and unless indicated the phase is x. Constant-time (T period) ^15^N Echo-Antiecho (EA) sampling is implemented with the aid of t1 and t2=t1*(T-4ζ)/(T+4ζ) delays and g1*EA1, g2*EA2, g3*EA3 and g4*EA4 gradients with classical EA1=EA2=(1,0.875), EA3=(0.6667, 1), EA4=(1, 0.6595) values; ^1^J_NH_ evolution delays were Δ = 2δ = 1/(4^1^J_CH_); constant time transverse relaxation delays, T are 40 and 60 ms; ζ delay vary from -T/4 to +T/4 modulating transverse relaxation from TROSY to antiTROSY under investigation; W_3_ is a 500 μs long both ^13^C^a^ and ^13^C° inversion adiabatic Chirp (Crp60,0.5,20.1 for 600MHz spectrometer) pulse with offset at 95 ppm. W_2_ is a 1500μs (for 600MHz spectrometer) long 180° Reburp pulse (offset at Hn center = 8.65ppm). W_1_ is a 1000 μs long 90° Sinc “down” water flip-back pulse (offset at 4.7 ppm). Gradient pulses with squared sine shape (SMSQ10.100) and 1 ms length except for g1 (0.5 ms) are used. The g1-g8 gradient strengths are as follows: 40%, 40%, 60%, 60.26%, 57%, 47%, 67%, 37% whereas 100% corresponds to ca 53 G/cm. The default phase is x and the phase cycle is: φ1 = 4(45°, -45°); φ2 = 90°; φ3 = 90°; φ4 = 4(0°, 180°); φ5 = 2(0°, 0°, 180°, 180°); φ6 = -90°; φ7 = (0°, 0°, 0°, 0°, 180°, 180°, 180°, 180°); φ8 = (180°, 180°, 180°, 180°, 0°, 0°, 0°, 0°); φrec = 4(0°, 180°). For each EA successive value φ2, φ3, φ6 and for t1 value φ1, φ4 and the phase of the receiver are incremented by 180° respectively. Similar to the original constant time experiment(Liu & Prestegard, 2008), the ^1^H signal position in all planes is the same and corresponds to TROSY peak, whereas ^15^N signal position ν^15^N-^1^J_NH_*ζ*2/T is a function of ζ delay (varying from –T/4 to +T/4).

Protein transcription was initiated by 0.5 mM isopropyl-β-D-1-thiogalactopyranoside (IPTG) to the culture after lowering the temperature to 20°C for overnight incubation. After centrifugation cells were resuspended in lysis buffer (20 mM Tris pH 7.5, 250mM NaCl, 20 mM Imidazole) supplemented with Complete protease inhibitor (Roche). Cells were lysed by sonication and cell debris was pelleted by centrifugation. The supernatant was loaded on a 5ml HisTrap FF column (Cytiva) and the protein was eluted with 500mM imidazole and concentrated down to 5ml with an Amicon Ultra centrifugal filter. After the purification via size exclusion chromatography with a HiLoad® 16/60 Superdex®, 75 PG, protein was eluted in 25mM NaPhosphate buffer, 100mM NaCl buffer and concentrated to 200mM.

### NMR experiments

NMR experiments were acquired on Bruker Avance III spectrometers operating at 14.1 T, corresponding to 600 MHz, equipped with a 5mm cryo-enhanced QCI-P probe. To improve relaxation parameters of the Psr_Sp_ the experiments were performed at 308 K temperatures. Backbone resonance assignments for Psr_Sp_ were obtained(Sandalova *et al*., 2024) and previously submitted by us to the BioMagResBank with accession code **52556**. Data were processed by TopSpin 4.06 (Bruker, Billerica, MA, USA), and analysed using CcpNmr2.4.2(Vranken *et al*., 2005) and Dynamics Center 2.8 (Bruker, Billerica, MA, USA).

### Pulse sequence for ^15^N-^1^H CSA/DD cross-correlation

A set of pulse experiments for measuring backbone ^15^N-^1^H chemical shift anisotropy/dipole-dipole (CSA/DD) cross-correlations in proteins, usually called H2, have been presented previously (Liu & Prestegard, 2008, Ferrage & Dorai, 2018). The scheme, where the ^15^N chemical shift evolution and modulation of signal intensities by cross-correlation are combined during a constant time period, was shown to provide superior signal-to-noise ratio (Liu & Prestegard, 2008). The new pulse sequence presented in **Figure 1** incorporates several features designed to minimize systematic errors in this approach: a) We found that application of ^15^N coding echo-anti echo gradients (g1 and g2) across all six intervals, where CSA/DD effects and sampling occur, minimizes the possible systematic errors of shaped and hard pulses with defocusing of residual undesired coherences. b) A classical ST2-PT TROSY block (Pervushin *et al*., 1998, Brutscher, 2024) with g1, g2, g3 and g4 echo-antiecho gradients and φ2 and φ3 phases are used for sampling and enhanced filtering of the TROSY component and simple preliminary selection through 2δ evolution. c) A ^1^H Reburp inversion pulse (W_2_), selective on amide protons, preserves water magnetization along the +Z axis and ensures uniform water (saturation) state for all ζ delays. d) W_2_ additionally ensures uniformity with respect to the ^2^J_NH-Hα_ scalar coupling evolution across all ζ delays. Note that a non-negligible fraction of Hα protons are typically present even in deuterated protein samples. e) All water flip-back pulses (W_1_) are placed outside the periods of the magnetization transfer over the ^1^J_NH_ coupling and are followed by gradients.

#### Determination of ^1^H-^15^N CSA/DD cross correlation (H2), relaxation rate R1 and ^1^H-^15^N Nuclear Overhauser Effect (NOE)

Backbone relaxation parameters R1 were recorded in pseudo 3D mode with randomized and interleaved (Korzhnev *et al*., 2001) relaxation delays using a standard Bruker pulse sequence, TROSY-version modified by Bax and colleagues in (Lakomek *et al*., 2012) trt1etf3gpsitc3d.3. Spectral widths (SW^1^H) of 16 ppm over 1024 complex points in the ^1^H dimension and spectral widths (SW^15^N) of 40 ppm and 128 complex points in the ^15^N dimension with 32 transients (NS) were used. R1 values were determined from a series of 11 relaxation delays: 10, 90, 192, 260, 380, 480, 690, 980, 1220, and 1444 ms. The errors in the R1 experiment were defined by an exponential decay fitting.

Backbone ^1^H-^15^N steady-state heteronuclear NOEs were measured using TROSY-type experiments (Zhu *et al*., 2000) implemented in Bruker pulse sequence, trnoeetf3gpsi3d.3. 2D experiments, including acquisition of NOE-enhanced and unsaturated spectra, were collected using D1 = 1s with a follow up ^1^H saturation time of 3s, spectral widths (SW^1^H) = 16 ppm and 1024 complex points in the ^1^H dimension, and (SW^15^N) = 40 ppm with 256 complex points, NS = 32. NOE values were obtained by dividing ^1^H-^15^N peak intensities in an NOE-enhanced spectrum by the corresponding intensities in an unsaturated spectrum, with an error of about 5% for all NOE experiments.

A complete set of ^1^H-^15^N CSA/DD cross correlation relaxation rates, H2, for the backbone amides were acquired at 600 MHz utilizing the pulse program presented above **Figure 1**. Experiments were performed using NS=24 on a time domain grid of 1 K x 128 complex points with spectral width/acquisition time of 15 ppm/114 ms for ^1^H and 40 ppm/53 ms for ^15^N dimensions with D1=1s and a constant time delay of T=0.06s. H2 values were determined from a series of 9 relaxation delays: - 0.05, -0.0375, -0.025, -0.0125, 0.0, 0.0125, 0.025, 0.0375, 0.05. Carrier positions: ^1^H, H_2_O frequency (4.698 ppm); ^13^C, 95 ppm; ^15^N, 118.0 ppm. Mirror image linear prediction was used for constant-time ^15^N sampling, doubling resolution without introducing exponential decay artefacts.

### Theoretical simulations: AlphaFold3 as a starting point for full atomic Molecular Dynamic

We utilized the Google DeepMind AlphaFold3 service [https://www.nature.com/articles/s41586-024-07487-w] to predict five Psr_Sp_ protein conformations. These predicted conformations did not account for solution properties such as pH values, temperature, ionic strength, etc, so there is a need to align the simulations to our experimental conditions. For Molecular Dynamics (MD), the charge of the protein residues was calculated at pH=6.8, with all histidine residues displaying neutral charges. The ionic strength was set to 138 mM, combining buffer and salt concentrations.

All MD simulations were performed using Gromacs version 2023.1 (Abraham *et al*., 2015) with the all-atom force field charmm36-mar2019_cufix.ff [https://github.com/intbio/gromacs_ff/tree/master/charmm36-mar2019_cufix.ff], including a refinement of the Lennard-Jones (Schwerdtfeger & Wales, 2024, Qiu *et al*., 2021) parameters (CUFIX) (Yoo & Aksimentiev, 2018). The protein was centred in a periodic cubic box (100Å), and Na^+^ (95) and Cl^-^ (83) ions were added to emulate the ionic strength and achieve electro-neutrality, as the protein had the total charge of (-12). Long-range electrostatic and van der Waals interactions were considered with a 10Å cut-off.

The predicted protein conformations underwent energy minimization to ensure a reasonable starting structure in terms of geometry and solvent orientation. Convergence was achieved at a maximum force of less than 1000 kJ/mol/nm for any atom. The potential energy was then used to select the best starting structure from the five AlphaFold3 predictions (Potential energy value for each structure: 1) -1.596e+06 kJ/mol 2) -1.562e+06 kJ/mol 3) -1.575e+06 kJ/mol 4) -1.608e+06 kJ/mol 5) -1.593e+06 kJ/mol). The fourth structure with the lowest potential energy was chosen for MD simulations.

Equilibration was conducted in two phases: the NVT (number of particles (N), volume (V), and temperature (T) are constant) ensemble for 100 ps, where the temperature of the system should reach a plateau at the desired value. Temperature was set to 308 K. And the NPT (number of particles (N), pressure (P), and temperature (T) are constant) ensemble for 100 ps until the system reached equilibrium, as indicated by a plateau in pressure and density values. A modified Berendsen-type (V-rescale) thermostat and a Parrinello-Rahman barostat were employed. Hydrogen-containing covalent bonds were constrained.

Following equilibration, MD simulations continued as a production run for 3000 ns under the same conditions. System stability was assessed using standard tools in Gromos (Abraham *et al*., 2015), monitoring temperature, pressure, energy, and periodicity. Visual inspection and RMSD analysis were performed in the xmgrace program (Turner PJ. XMGRACE, Version 5.1.19. Center for Coastal and Land-Margin Research, Oregon Graduate Institute of Science and Technology, Beaverton, OR; 2005).

Cluster analysis were performed for trajectory regions 700-1200 ns and 1750-2250 ns. 1000 trajectory frames with 500 ps steps for each trajectory region were partitioned into 2 arrays of 10 clusters each using the Gromos algorithm.

### Individual MD trajectory analyses with back-calculation of theoretical ^15^N relaxation parameters

MD trajectory analyses with the backbone ^1^H-^15^N vector extraction and approximation of autocorrelation function C(t) with the subsequent spectral density function J(ω) calculations were utilized as previously described (Agback *et al*., 2023), with the aid of the “Mathematica” software package ***<u>[Wolfram Research]</u>*** and the MD Analysis external library [mdanalysis.org]. Back-calculation of classical NMR ^15^N relaxation parameters R1, H2 and NOE as a function of J(ω) were also performed as previously described (Agback *et al*., 2023, Showalter & Bruschweiler, 2007), whereas ^1^H-^15^N CSA/DD cross-correlation contribution to transverse relaxation, denoted as:

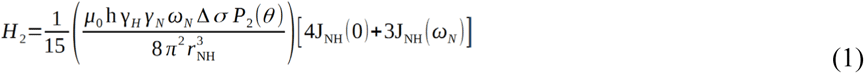

where the spectral density function was: 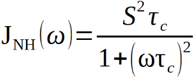

Where µ*_o_* is the vacuum permeability; h is Planck’s constant; *Y_H_* and *Y_N_* are the gyromagnetic ratios of ^1^H and ^15^N respectively; Δσ is the chemical shift anisotropy (CSA) of ^15^N with CSA=-166ppm (Robson *et al*., 2021); *r_NH_* = 1.023Å (Yao *et al*., 2008); *ω_N_* is the Larmor frequency of ^15^N at 600 MHz; P_2_(θ) is the Legendr 2^nd^ degree polynomial; θ=17° is the CSA/NH vector angle (Chill *et al*., 2006); J(ω) is the NH auto-correlation spectral density function.

## RESULT

### Selection of relaxation parameters for MD trajectory verification: comparison of R2 and H2 relaxation data

In the method presented here to verify MD trajectories and their segments, experimental criteria are necessary to quantitatively compare both the amplitudes (S^2^ order parameter) in R2 and H2 (**Figures** 2c, d) and relaxation times, (τ_e_), in R1 and NOE (**Figures** 2a, b) of intramolecular motions in proteins. A key criterion for the selection of dynamic parameters for measurement and back-calculation is the minimization or elimination of systematic errors in the experimental procedures.

**Figure 2:**
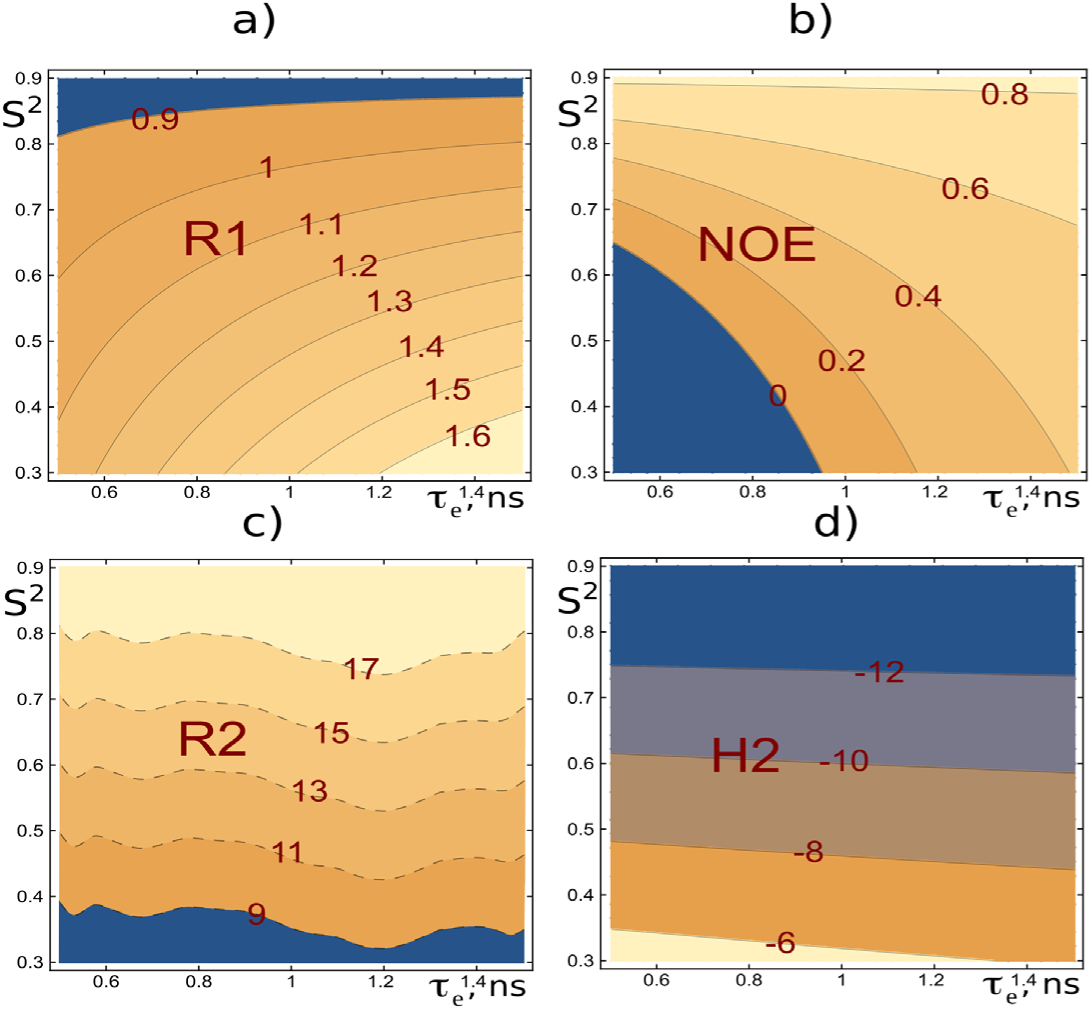
Schematic profiles of relaxation parameters. R1, NOE, R2 and H2 are shown in panels (**a**), (**b**), (**c**), and (**d**), respectively as functions of the internal motion parameters τ_e_ and S^2^=S^2^_fast_*S^2^_slow_ (with S^2^_fast_ fixed at 0.9) for a protein with an overall correlation time τ_c_ of 14.7 ns. These profiles were generated using the classical Extended Lipari-Szabo model, which utilizes three internal motion parameters (τ_e_, S^2^_fast_, S^2^_slow_ and the overall correlation time τ_c_ (Lipari & Szabo, 1982b, a, Chevelkov *et al*., 2007, Korzhnev *et al*., 2001). The plots illustrate the most typical ranges for S^2^ and τ_e_. The dashed profiles in the R2 panel (c) artistically depict the influence of systematic experimental errors, such as Rexp, on the measurements.

Typically, the set of relaxation parameters used to characterize protein backbone dynamics includes R1 (**Figure** 2a), NOE (**Figure** 2b), and R2 (**Figure** 2c) (Showalter & Bruschweiler, 2007). However, the practical use of R2 relaxation rates is hindered by several systematic errors. The most significant issues are: (i) the contribution of chemical exchange (Rex), (ii) relaxation delay-dependent modulation of signal intensities by the water saturation, caused by the direct exchange of amide protons and cross-relaxation-mediated coupling between protein and water magnetization, and (iii) off-resonance effects in CPMG experiments (Korzhnev *et al*., 2000).

As alternative to R2 one can use H2 experiment (Kämpf *et al*., 2018). In this study, we use carefully calibrated measurements of H2, which are free from the aforementioned issues, to verify the amplitudes of angular motions of the intramolecular NH vectors motions.

In **Figure S1(a)**, the first 2D plane at delay ζ equal to 0 in the H2 experiment for the Psr_Sp_ protein is presented, with the experiment performed at a constant time interval of T = 60 ms. A comparison of the first increment of the H2 experiment (**Figure S1(e)**) with (black curve) and without (red dashed curve) water flip-back selective pulses show that implementing these pulses in the pulse sequence (**Figure 1**) prevents saturation of amino proton signals by water and improves the intensities of the observed signals by more than 1.5 times.

At constant time delay T=60, the intensity of the signal, shown in the projection through the 306G cross-peak, decreases approximately 2.5-fold, as evidenced by the comparison of the black curve (first 2D plane) and the red dashed curve (last 2D plane) in **Figure S1 (f)** for ζ delays of 0 and 0.098 s, respectively.

As an example, to illustrate the quality of the experimental data obtained under the aforementioned conditions, the intensity changes as a function of delay ζ, fitted to exponential curves to determine the H2 relaxation rates for the selected residues 174G, 307S, and 306G, are shown in **Figures S1(b)**, **(c)**, and **(d)**, respectively.

The final experimentally obtained H2 relaxation data, along with their fitted errors for the PsrSp protein, are presented in **Table S1**, and used for the validation of the MD-derived conformation trajectories in **Figure 4 (b)**.

### Determination of the isotropic rotational tumbling time of protein

The classical approach (Cavanagh *et al*., 2007) for estimating rotational tumbling time (τ_c_) utilizes R2 and R1 values. However, as mentioned above, R2 values are affected by residual R_ex_ contribution, CPMG off-resonance effects and several other artifacts. To address these issues in estimating τ_c_ values, we used H2 values (Equation 1) instead of R2. The Lipari-Szabo model (Korzhnev *et al*., 2001) was applied to stable backbone NH vectors with minimal R1 and H2 experimental errors, providing two key values: S^2^ and τ_c_ for all NH groups by numerically solving Equations (1) and (2)

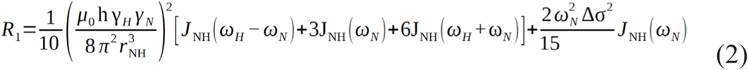

Using this approach, the τ_c_ value for the Psr_Sp_ protein was estimated by selecting amino acids with S^2^ > 0.8, resulting in 76 stable NH groups. The mean τ_c_ value was found to be 14.7 ± 0.2 ns, supporting the predominantly monomeric form of Psr_Sp_ in solution. This experimental τ_c_ value is consistent with predictions made using an empirical equation (Cavanagh *et al*., 2007) that accounts for temperature and molecular weight. At 35°C, the predicted *τ*_C_ values were 15.6, 31.1 and 46.6ns for the monomeric, dimeric and trimeric forms of Psr_Sp_, respectively.

### Identification and Validation of Structural Ensembles of the Psr_Sp_ protein Based on backbone relaxation dynamic

We have recently determined the crystal structure of the extracellular region of the *Streptococcus pneumoniae*-associated polyisoprenyl-teichoic acid-peptidoglycan teichoic acid transferase Psr_Sp_ (residues 131-424) (Sandalova *et al*., 2024), whose topology diagram is presented in **Figure 3(b)**. Based on the near-complete assignment of the ^1^H, ^13^C, and ^15^N backbone resonances, as well as the ^13^Cβ side chain resonances for the Psr_Sp_ domain (for amino acid sequence of the Psr_Sp_ construct **Figure 3(a)**), the secondary structure of Psr_Sp_ in solution was determined and compared with the high-resolution X-ray crystal structure. Additionally, dynamical S^2^ predicted order parameters were extracted and compared with structural information and the crystallographic B-factor allowing us to qualitatively evaluate the flexibility of this protein. Although the assignment of the backbone resonances is an essential first step, this experimental data alone is insufficient to extract the conformational ensembles reflecting the flexibility of the protein.

**Figure 3.**
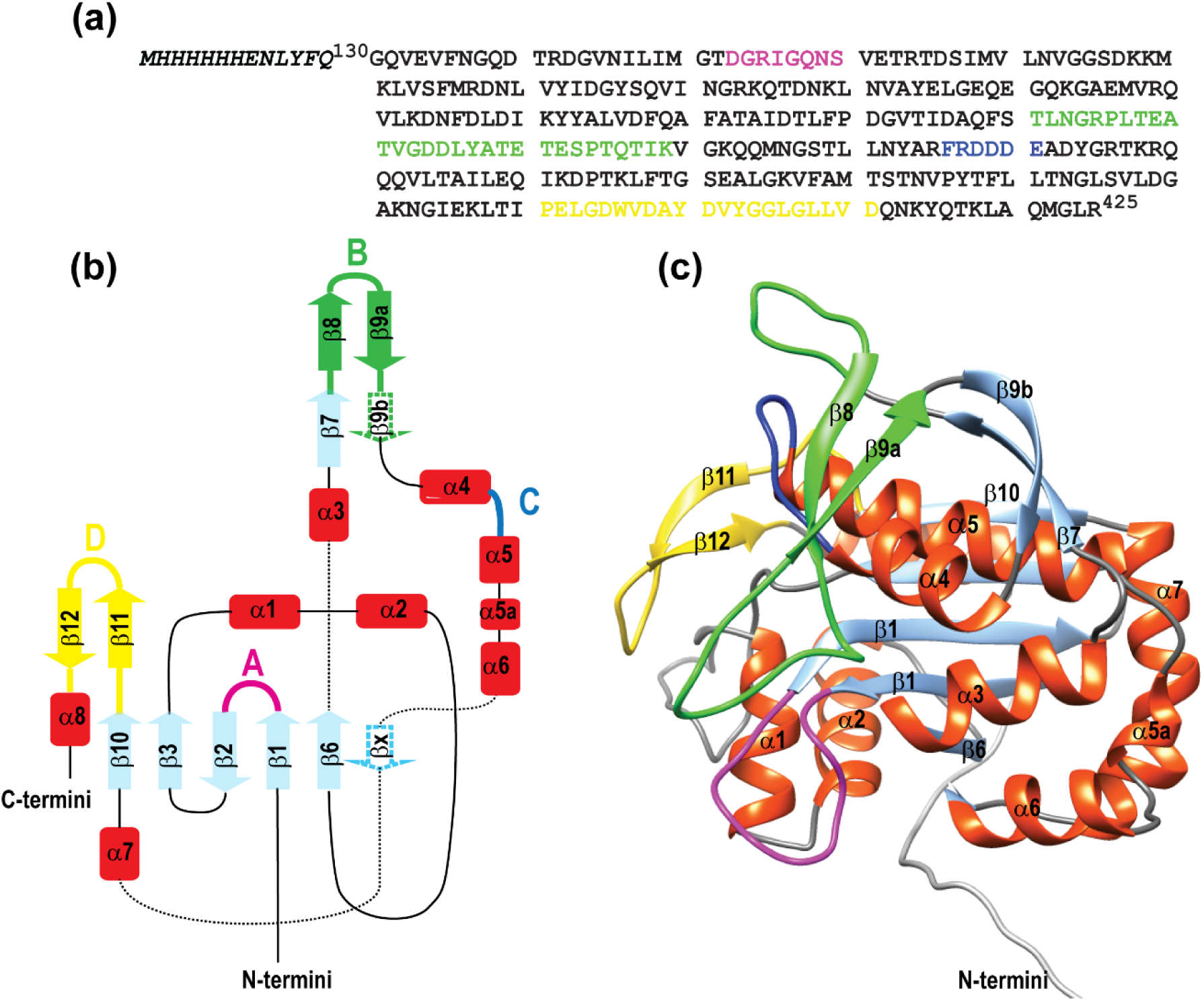
Topology diagram of Psr_Sp_ protein. (**a**) Amino acid sequence of the Psr_Sp_ construct. The TAG aa are indicated in italic. The topology diagram of Psr_Sp_ is shown on the (**b**) left Panel while cartoons representation of the AlphaFold 3 molecular model of the extracellular region of Psr_Sp_ are shown on the right **(c**) Panel colored according to the topology diagram. As for coloring, all α -helixes are red, β-strand are light blue, except β11, β12 which are yellow, and β8, β9 a,b green; linker are black; except for loops and linkers belonging to the four regions A-D that are suggested to be important for substrate binding to Psr_Sp_ are colored in pink, green, dark blue and yellow, respectively.

To comprehensively explore the conformational space and generate an ensemble of Psr_Sp_ conformations, we employed a strategy previously developed by our team (Agback *et al*., 2023). This approach leverages molecular dynamics (MD) trajectories validated against experimentally determined relaxation parameters of the NH backbone. Notably, in this protocol, the H2 relaxation parameter was used in place of R2 to enhance accuracy. In comparison to our previously established protocols (Agback *et al*., 2023) which utilized NOE-based NMR and X-ray structures as starting points, the present study explores the validation of a novel approach where an AlphaFold-generated structure serves as the starting structure for free MD simulations. The motivation for testing this method lies in the accessibility of AlphaFold (AF), which can generate both single molecular models and conformational ensembles (Wallerstein *et al*., 2024, del Alamo, DeSousa, *et al.*, 2022, del Alamo, Sala, *et al.*, 2022, Jumper *et al*., 2021), offering a faster and more cost-effective alternative to traditional NMR or X-ray structures for initiating MD simulations. This approach enables the generation of trajectory intervals used to back-calculate the relaxation parameters R1, H2, and NOE for the Psr_Sp_ ^1^H-^15^N amide backbone. The AF-predicted three-dimensional structure of the Psr_Sp_ protein, used as the starting point for a 3 μs free MD simulation, is presented in **Figure 3(c)**. In particular, the root mean square deviation (RMSD) between the AF structure used in this study and the experimentally determined X-ray structure of Psr_Sp_ (Sandalova *et al*., 2024) is 1.32 Å for 277 aligned residues. The N-terminal 14 residues, loop A, and approximately 10 residues following helix 5 **Figure 3(c)** are excluded from alignment due to significant positional deviations.

After 700 ns of free MD simulation equilibration, the RMSD of backbone heavy atoms for dynamically stable residues of Psr_Sp_ went through three plateaus (**Supplementary Figure S2**). Three distinct 500 ns time intervals were selected for back-calculation of the relaxation parameters R1, H2, NOE and S^2^ of the Psr_Sp_ protein, and defines ensembles I, II, and III, which correspond to trajectories from 700–1200 ns, 1750–2250 ns, and 2500–3000 ns, respectively. Segmentation of MD trajectory for back calculation of the relax parameter were also used in IDP studies (Kämpf *et al*., 2018).

The back-calculated theoretical versus experimental ^1^H-^15^N R1, H2, and NOE parameters for the Psr_Sp_ backbone are shown for all three trajectories in **Figure 4**. The secondary structure elements based on the crystal structure of Psr_Sp_ (Sandalova *et al*., 2024) are presented at the top of **Figure 4**.

**Figure 4.**
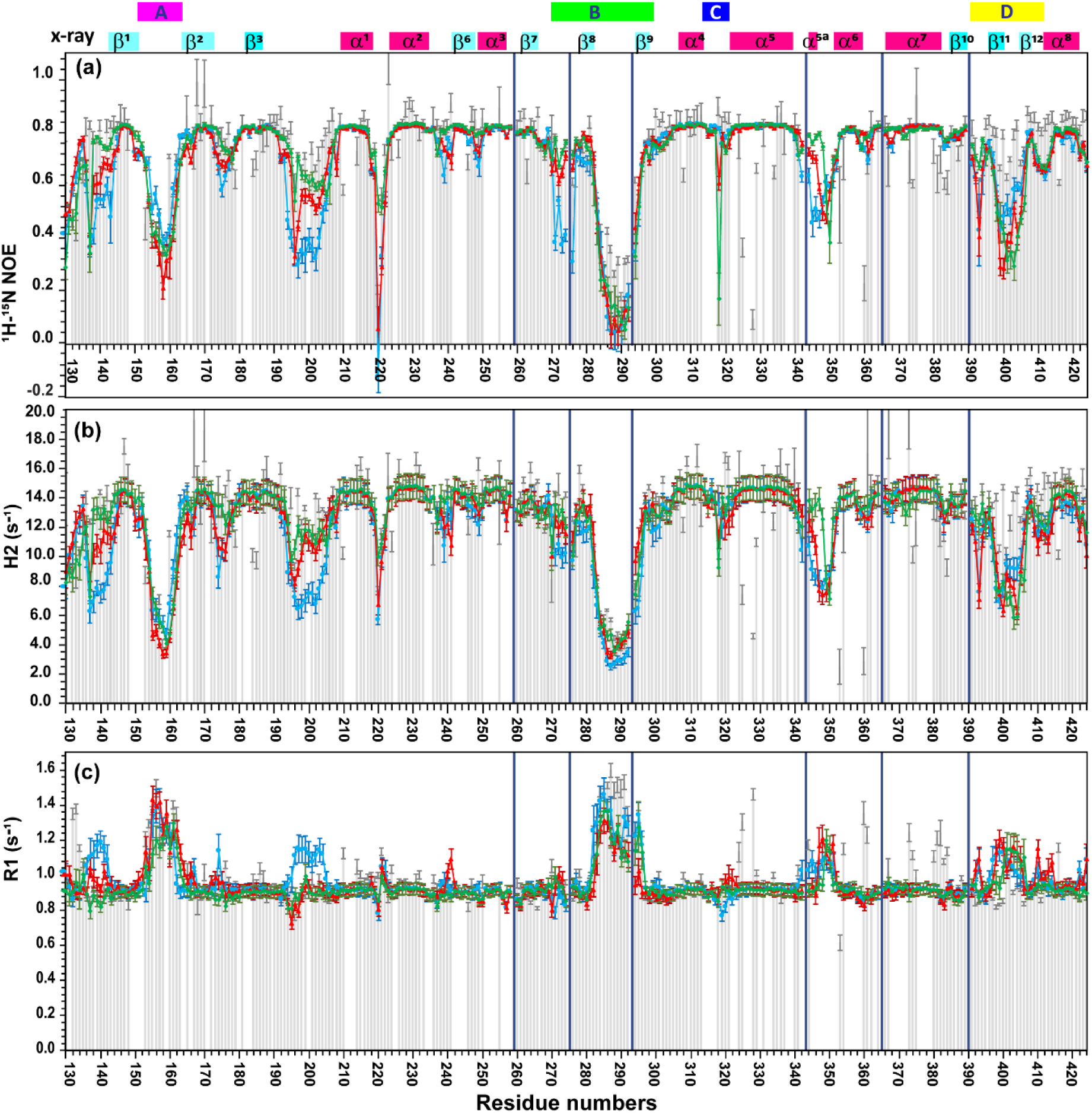
Psr_Sp_ amide backbone, ^15^N(H), dynamic parameters obtained on 600MHz spectrometers. The relaxation parameters (**a**) heteronuclear ^1^H-^15^N NOE values; (**b**) the transverse relaxation time H2(s^-1^) (**c**) longitudinal relaxation rate R1(s^-1^) The experimentally obtained R1(s^-1^), H2(s^-1^) and NOE values are presented by light grey solid bars. The theoretically predicted dynamic parameters R1, H2 and NOE, obtained from three trajectories, are displayed by solid lines and coloured in dark green trajectory (700-1200), red trajectory (1750-2250) and light blue trajectory (2500-3000ns) for ensembles I, II and III, respectively. The error bars obtained by the bootstrapping analysis are shown at the level of one s from the predicted parameters. Long black bars show the position of the proline amino acids. The secondary structural elements of the Psr_Sp_ are shown at the top of the panels, based on the crystal structure of Psr_Sp_ proteins obtained previously (Sandalova *et al*., 2024).

The dynamic parameters ^15^N R1, H2, and NOE parameters show good agreement across all three back-calculated trajectories and with experimental data for the most stable secondary Psr_Sp_ structure elements determined by X-ray crystallography (Sandalova *et al*., 2024). As shown in **Figure 4**, this is evident for the β1-β10 beta strand and the α1, α2, α4 and α8 helices, where the differences between the calculated curves and experimental data fall within the error margin. This consistency indicates absence of the systematic shifts between theoretical and experimental results, validating the back-calculation protocol presented in this study.

Unfortunately, due to lack or high errors of the experimental data, validating back-calculated trajectories I-III against NMR experimental data was not possible for the α3, α5 and α7 helices. As reported in our previous work (Sandalova *et al*., 2024), the resonances from these amide protons were not observed because of exceptionally slow deuterium-to-proton exchange. Importantly, the back-calculated trajectories I-III closely align with the X-ray structure of Psr_Sp_, particularly for the sequence regions corresponding to the α3, α5 and α7 helices.

There is a consensus among the ^15^N R1, H2, and NOE values and the experimental data in two regions of interest, A and B (Figure 3b), which demonstrate high amide mobility in the loops regions. This corroborates the observation that the NOE values drop from approximately 0.8 to 0.2 and the H2 rates decrease from 14 to around 3 s ^-1^.

Important discrepancies are observed between the H2 values of trajectories I–III and the experimental data in regions A and B (**Figure 4(b)).** As previously noted, the experimental H2 relaxation data exclude contributions from slow exchange Rex, which is expected in the loops of regions A and B. Detailed examination of these regions reveals that trajectory I aligns more accurately with the experimental H2 data, even though trajectory II also shows reasonable proximity to the experimental values. The main differences in the ^15^N R1, H2, and NOE parameters calculated for trajectories 1-III are observed in the residue ranges 190–210, 135–145 and 265-275. Among these, trajectory I provides the best fit to the experimental data, while trajectory III shows a poor fit and cannot be considered as verified. In conclusion, the most important result is that differences between two consecutive (**Figures S1**) trajectories, I and II are revealed by the differences in ^15^N R1, H2, and NOE relaxation parameters.

### Determination of S^2^ order parameters from trajectories

The theoretical order parameters S^2^ for the backbone of Psr_Sp_ were back-calculated from the corresponding MD trajectory as a value of the backbone ^1^H-^15^N vector autocorrelation function C(t) at five times the rotational correlation time 5×τ_c_=5×14.7 ns, as described by us earlier (Agback *et al*., 2023). The ^15^N(H) order parameters (S^2^) extracted from the theoretical relaxation profiles of ensembles I-III are presented in **Figure 5**.

**Figure 5.**
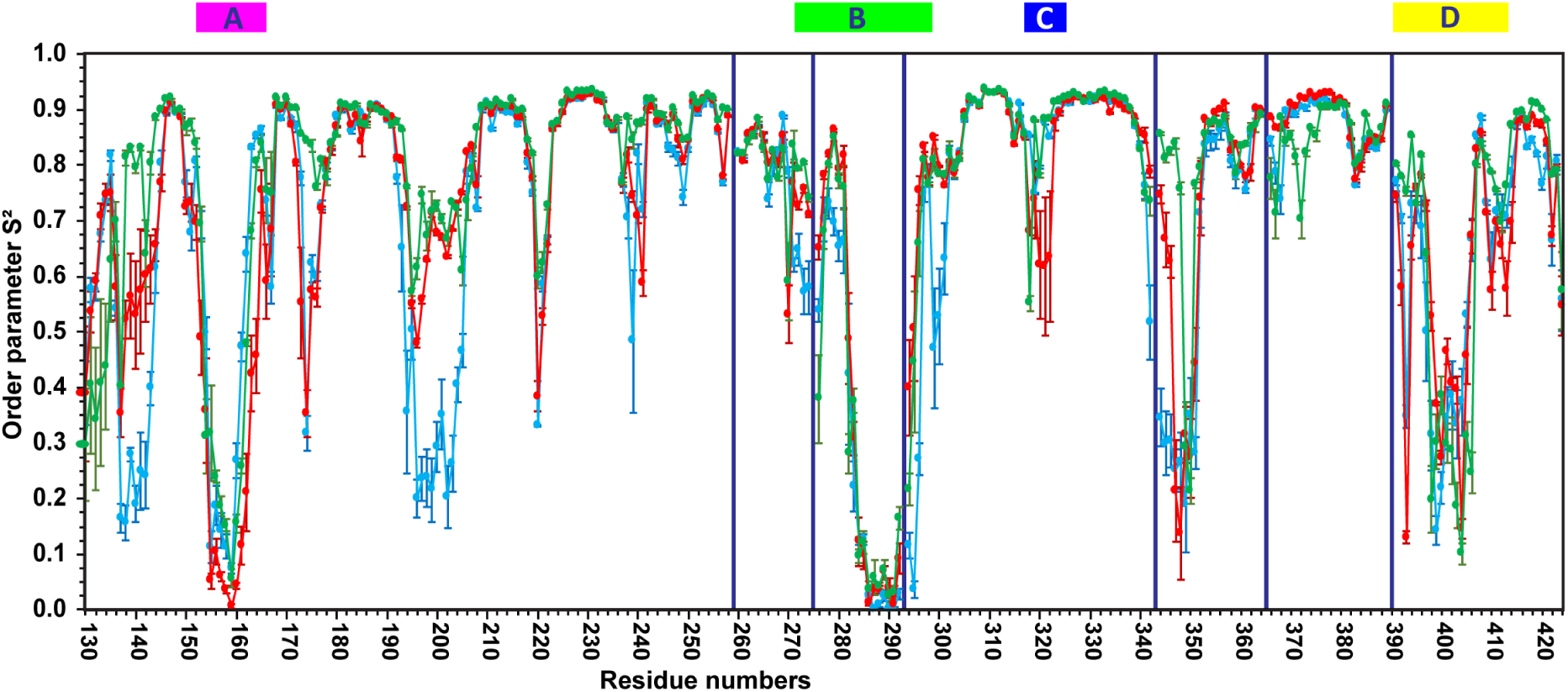
Order parameters S^2^ for Psr_Sp_ amide backbone ^15^N(H), calculated from MD trajectories are shown in green, red and blue for trajectories I (700-1200ns), II (1750-2250ns) and III (2500-3000ns). The order parameters were back-calculated from the corresponding MD trajectory as the value of the backbone ^1^H-^15^N vector autocorrelation function C(t) at 5×τ_c_=5×14.7 ns.

The extracted S^2^ parameters in regions A and B for all three trajectories (I-III) are very low, indicating high flexibility (**Figure 5** and **Figure 6(a)**). The S^2^ parameters reveal significant discrepancies between trajectories I and II compared to trajectory III. As shown in **Figure 5**, the latter exhibits greater flexibility in regions spanning residues 135-143, 190-210, 265-275, and 295-300. As shown above, trajectory III has the worst fit to the experimental relaxation parameters. For trajectories I and II, which show the best agreement between back-calculated relaxation parameters and corresponding experimental data, S^2^ values generally fall within the range of error.

**Figure 6.**
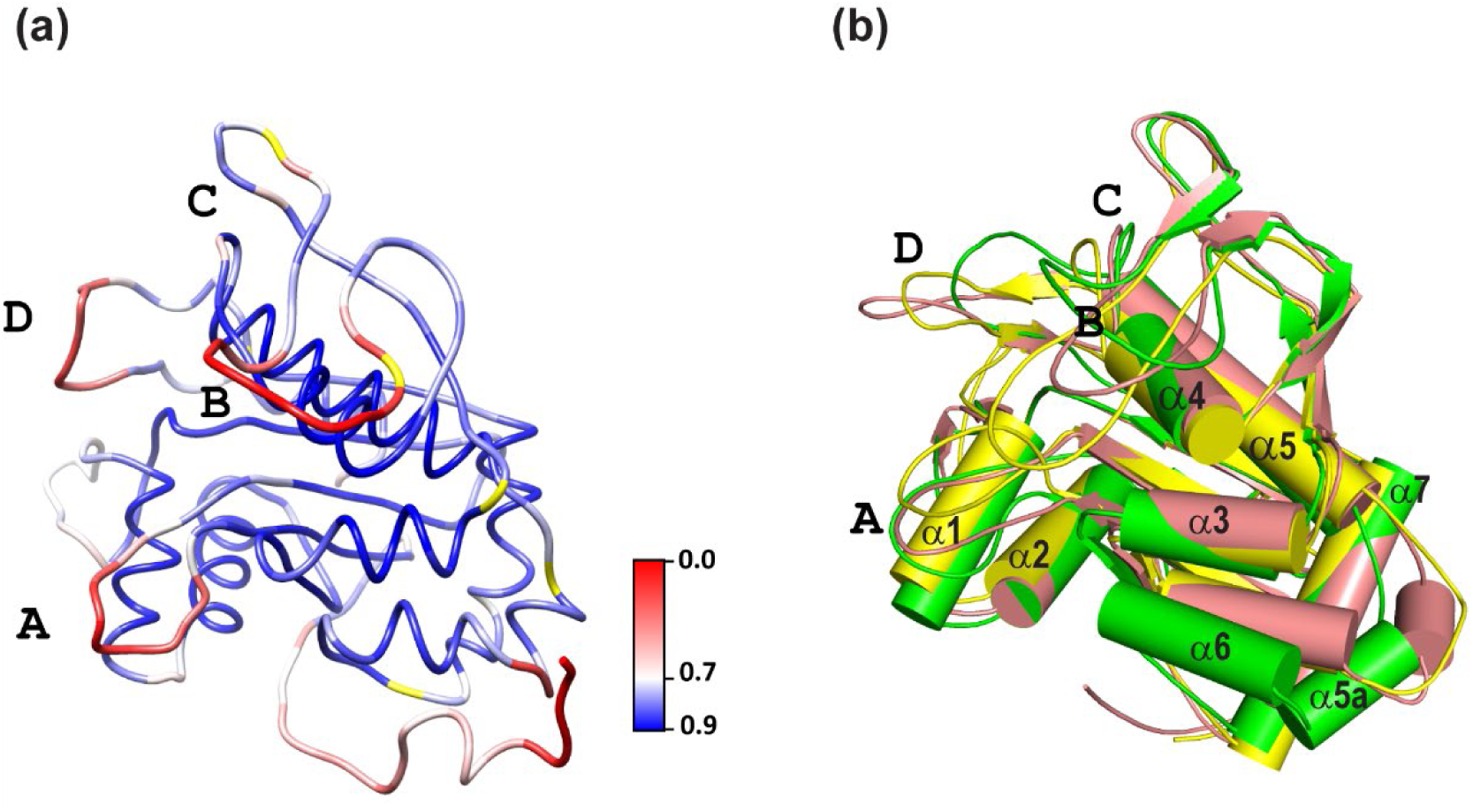
Psr_Sp_ amide backbone, ^15^N(H), order parameters S^2^ structural-dynamic models. (a) The distribution of the order parameters S^2^ ranging from S^2^=0.0 (red) to S^2^=1.0 (blue) is shown on the most representative structure obtained in cluster analysis of trajectories 700-1200ns. Proline and amino acids where S^2^ was not defined are shown by yellow. (b) Superposition of the most representative models of Psr_Sp_ from the trajectories 700-1200ns (green), 1750-2250 (pink) and 2500-3000 ns (yellow) demonstrates the mobility of α-helix 6. The main difference between these clusters, except the conformations of several long loops is the region between helices α5 and α7: a small additional helix α5a presented in the structure of trajectory 700-1200ns (green) while helixes α6 and α5a shift by 3-5Å in the structure of trajectory 1750-2250 (pink) and in the structure of 2500-3000 ns (yellow) instead of helix α5a a flexible loop is observed. Four regions A-D that are suggested to be important for substrate binding to Psr_Sp_ are indicated by capital letters, respectively.

In trajectory I, S^2^ values are consistently higher compared to trajectory II, particularly in the regions 135-143, 170-175, and 340-355. The later interval, 340-355, is especially noteworthy. In this region, trajectory I exhibits two stable segments in Psr_Sp_, with S^2^ values around 0.8 between residues 345-349 and 350-356. These segments correspond to the α5a and α6 helices, separated by a sharp flexible turn. The helices are depicted as green cylinders in **Figure 6(b)**, which also illustrates the superposition of the most representative models of Psr_Sp_ from trajectories I (700-1200ns green), II (1750-2250 pink) and III (2500-3000 ns yellow).

In trajectory II, the S^2^ values in the discussed region (345-349) decrease to 0.5-0.6, and drop further to 0.3-0.1 in trajectory III. This decrease corresponds to structural rearrangements between the α5a and α6 helices. Specifically, in trajectory II, the α5a helix shortens into a turn, and both α6 and α5a shift by 3-5Å relatively to their positions in trajectory I (700-1200 ns). In trajectory III, the α5a region further transforms into a flexible loop. These structural changes are depicted in **Figure 6(b)** by pink and yellow cylinders, respectively.

Cluster analyses were performed on free MD simulations of the two trajectories I and II, which were verified by back calculation of relaxation parameters as discussed here above. The populations of 10 clusters presented in **Table S2**. In **Figure S3,** for the two trajectories, are shown 10 representative structures, one for each cluster.

## Discussion

One of the most groundbreaking advances in structural biology over the past decade has been the recognition of the pivotal role that conformational heterogeneity plays in key biological processes. Recent reviews have highlighted diverse systems and mechanisms that underscore the significance of dynamic conformational ensembles (Nussinov, Liu, *et al.*, 2023b, Ramelot *et al*., 2023, Gampp *et al*., 2024). As a result, current research efforts are focused on several key directions within this emerging paradigm. Hotspots, often described as conserved regions within dynamic protein structures, act as “islands of stability” and play a critical role in binding energy and functional specificity (Bogan & Thorn, 1998, Kuttner & Engel, 2012). While traditionally regarded as structurally stable regions, increasing evidence underscores the importance of understanding the full conformational ensemble to accurately interpret hotspot behaviour (Zhuravleva *et al*., 2004, Lexa & Carlson, 2011, Alvarez-Garcia & Barril, 2014)

Protein flexibility, particularly in ligand binding (Zerbe *et al*., 2012, Rosell & Fernandez-Recio, 2018, Yu *et al*., 2022, Zhang *et al*., 2022, Paquete-Ferreira *et al*., 2024) is vital for modulating activity through mechanisms such as shifts in conformational balance, formation of ligand-induced conformations, and active site mobility required for allosteric signalling (Nussinov, Zhang, *et al.*, 2023). These findings highlight the limitations of static models in capturing dynamic interactions and underscore the necessity of adopting dynamic perspectives in structural biology (Tsai *et al*., 2008, Lexa & Carlson, 2011, Nussinov, 2016). Efforts to generate accurate protein conformational ensembles have led to the development of advanced methodologies. Cryo-EM, while provides conformation ensembles of large protein complexes and biological machines, often suffers from resolution limitations. All-atom molecular dynamics simulations have significantly advanced the theoretical exploration of dynamic ensembles for biologically relevant systems (Dauber-Osguthorpe & Hagler, 2019, Schlick & Portillo-Ledesma, 2021, Schlick *et al*., 2021, Zhang *et al*., 2022). Meanwhile, AlphaFold can predict structural ensembles, although the biological relevance of these ensembles requires further validation (del Alamo, DeSousa, *et al.*, 2022, del Alamo, Sala, *et al.*, 2022, Wallerstein *et al*., 2024). Despite these advancements, a critical challenge remains: experimentally validating these theoretical conformational ensembles.

### Experimental Validation of Conformational Ensembles

NMR spectroscopy, particularly residual dipolar coupling (RDC)(Robertson *et al*., 2021), NOE-based and chemical shift (CS) methods, offers potential solutions for validation of conformation ensembles. Conformers obtained through AlphaFold, Cryo-EM, or MD trajectories can be validated by back-calculating the chemical shifts of backbone and side-chain nuclei, and comparing these with experimentally determined values. Software, such as ShiftX2 v.1.10 (Han *et al*., 2011), provide a comprehensive chemical shift prediction package based on PDB structures, enabling the assessment of structural ensembles through these comparisons. However, the relatively low accuracy of existing chemical shift prediction methods limits the reliability of this approach for discriminating between different conformational ensembles. Protein structures obtained by traditional NOE-based NMR method generally align well with X-ray crystallography data. Careful quantitative use of ^1^H−^1^H NOEs has been employed to generation structural ensembles of small sized protein systems (Lomize *et al*., 1992, Vögeli *et al*., 2012, Vögeli *et al*., 2016, Wenchel *et al*., 2024) However, large, deuterated proteins and intrinsically disordered proteins (IDPs) pose unique challenges due to their flexible and heterogeneous landscapes. While NOE-based conformational ensembles provide valuable insights into protein dynamics, they come with notable limitations.

A key distinction between these methods lies in their sensitivity to conformational variability and the nature of the parameters being measured. The use of ^1^H−^1^H proton cross-relaxation (^1^H−^1^H NOE) as a criterion for constructing a network of intramolecular contacts often results in conformational bias and degeneracy because of the short-range nature of the ^1^H−^1^H NOE interactions rapidly diminishing with the proton-proton distance (R^−6^ dependency). This limitation hinders the experimental derivation of reliable ^1^H−^1^H NOE based conformational ensemble.

Furthermore, back-calculations of ^1^H−^1^H NOE to validate all-atom MD trajectories remain challenging. Unlike ^1^H−^15^N relaxation, which is described primarily by local dipole-dipole interactions and chemical shift anisotropy (CSA), mainly reflecting fluctuations in orientation the ^1^H−^1^N vector, the ^1^H−^1^H NOE is also affected by the spin-diffusion and fluctuations in inter-proton distances. Moreover, measured ^1^H−^1^H NOE values are influenced by slow conformational exchange and exchange of labile protons with water, making them poorly suited for validating dynamic ensembles.

In practical applications NOE values represent time-averaged conformational structure, which fail to capture temporal fluctuations in protein structure. However, the aim of our study, is to establish a quantitative rather than qualitative criterion for this purpose.

### Relaxation-Based Validation of Conformational Ensembles

As an alternative, in our recent work on the NS3_pro_-NS2B protein_(Agback *et al*., 2023), we demonstrated that the combination of MD simulations with NOE-derived restraints led to poor agreement with relaxation experimental data, resulting in inaccurate conformational ensembles. To address these limitations, we introduced a relaxation filter approach that integrates experimental relaxation measurements with unconstrained MD simulation and back-calculations of relaxation parameters.

In the present study, we show that CSA/DD cross-correlation relaxation rates, H2, which, unlike R2, are free from contributions of in millisecond time scale conformational exchange and avoid other experimental artefacts, fit well with back-calculated relaxation data. We also present an improved version of the pulse sequence for measuring H2, which is sensitive, free from water saturation effects on the amide proton, and optimized for large proteins. The AF-free-MD-NMR-based method proposed in this study enables robust analysis of protein dynamics and experimental validation of the conformational ensembles.

Based on the crystallographic data from our earlier studies, we highlighted the highly dynamic nature of the extracellular region of the *Streptococcus pneumoniae* protein Psr_Sp_ (Sandalova *et al*., 2024). In this study, Psr_Sp_ (residues 131-424) was used as a model system to demonstrate the described above approach for obtaining and validating conformational ensembles. We report, for the first time, the relaxation dynamic parameters R1, NOE, and H2 for PsrSp (**Table S1** and **Figure 4**). These experimental data provide insights into the dynamic behaviour of PsrSp in its relaxed, ligand-free state. First, we determined the overall correlation time of the protein and confirmed that PsrSp behaves as a monomer in solution. This result aligns well with our X-ray crystallography data (Sandalova 2024). The next step involved back-calculating the R1, NOE, and H2 relaxation parameters for PsrSp to identify the conformational ensembles obtained from free MD simulations that best align with the experimental data. In our previous study (Agback *et al*., 2023), we used the X-ray structure of the Dengue II NS3-NS2B enzyme and three NOE-refined NMR structures as starting points. Here, we explored an alternative approach by using a single AlphaFold-generated structure of PsrSp as the starting point, and conducting one long free MD simulations. To our knowledge, our approach for the first time provides experimentally verified conformational ensembles based only on measurements of the R1, H2 and NOE relaxations without using any other additional experimental data.

Although for PsrSp we identified stretches of the MD trajectory that fit well with the relaxation data, in other cases it may happen that none of the ensembles generated in a long trajectory fits the experiment. This could be due to inappropriate starting structures, simulation conditions, or force fields, leading to incorrect sampling of the conformational space. Such MD trajectories should be discarded, and new ones produced to address these issues. For example, it is possible to start the simulation from another structure offered by AlphaFold. Nonetheless, if a conformational ensemble matches the experimental relaxation data, it may be considered as a plausible experimentally verified solutions.

### Identification of Conformational Ensembles

In this work, three distinct conformational ensembles (I-III) with varying population distributions were obtained from a single relatively long 3 µs MD simulation. The subsequent comparison of the relaxation parameters back-calculated from these ensembles with the experimental values (**Figure 4**), revealed that ensemble I, corresponding to the trajectory segment between 700–1200 ns, provided the best fit to the experimental data. The differences between ensembles I and II were primarily reflected in the rearrangement of the α6 and α5a helices (**Figure 6**), and in the population distributions of the 10 clusters derived from the trajectories (**Table S2**). Notably, flexibility in this regions was suggested in the crystal structure of Psr_Sp_ where of residues 342-348 adopted different conformations in three monomers present in the asymmetric unit of the crystal(Sandalova *et al*., 2024). Furthermore, residues corresponding to helix α6 are not observed in the electron density maps of all five available crystal structures of the homologous TagT protein from *Bacillus subtilis* (Schaefer *et al*., 2018, Eberhardt *et al*., 2012, Li *et al*., 2020). This points to the mobility of this region and potentially hints at its functional significance.

### Experimental Validation and Functional Implications

Figure 5 highlights the high sensitivity of this approach, as even subtle population shifts can be captured in the recalculated order parameters (S^2^). These findings also emphasize the importance experimental validation of the MD trajectories. For instance, we observed in this study that trajectory segment I (700– 1200 ns) provided the best fit to experimental data, whereas trajectory III (2500–3000 ns) was identified as the most significant outlier. S^2^ extracted from the trajectory I (Figure 5) corroborate our earlier findings(Sandalova *et al*., 2024) whereas trajectory III predicts flexibility in regions that are not supported by the new relaxation experiment and published crystallographic data. In trajectory I, regions A and B of Psr_Sp_ exhibit significantly higher flexibility than other parts of this key LCP protein. We speculate that this flexibility in loops A and B is functionally significant, potentially enabling Psr_Sp_ to catalyse the attachment of a wide range of glycopolymers to peptidoglycan.

### Deposition of Data and Structures

Finally, cluster analysis of the free MD simulation trajectories yielded 10 final structures for each of the two conformational ensembles (I and II) (**Figure S3**). These structures of trajectory I have been deposited in the Protein Data Bank (PDB) (Entry ID: D_4-G0H2) along with their S^2^ parameters, population values and their R1, H2 and NOE relaxations data experimentally obtained and back calculated from free MD simulation in the Biological Magnetic Resonance Data Bank (BMRB 52556). These data provide a valuable resource for further research, including the development of novel antibiotics targeting this essential protein.

### Conclusion

Conformational heterogeneity plays a crucial role in protein function, necessitating approaches that account for dynamic ensembles. Our study demonstrates the utility of combining free MD simulations with relaxation-based experimental data for validating the resulting conformational ensembles. Using only an AlphaFold-generated structure of Psr_Sp_ as the starting point and a set of backbone relaxation measurements, we identified biologically relevant dynamical conformational ensembles. These findings underscore the importance of experimental validation of MD trajectories. We demonstrated that only certain segments of the MD trajectory align well with experimental relaxation data, while others deviate significantly.

Additionally, we identified two flexible regions in Psr_Sp_, which may be involved in the catalytic function. The validated ensembles provide a valuable resource for further structural and functional investigation, including antibiotic development targeting this protein.

Finally, we note that the new approach presented in this study represents a powerful m for the reliable determination of protein dynamical conformational ensembles with minimal resources.

## AUTHOR CONTRIBUTIONS

BMS has contributed with production of all necessary protein variants, including their purifications, and developing a construct of labeled Psr_Sp_ proteins. DL, VO and TA wrote the original manuscript draft. DL, AA, VO and PA contributed with final editing of the manuscript. DL contributed to the development of the H2 pulse sequence and relaxation parameters back calculation. KR contributed with the MD simulations and trajectory analysis. TA and PA performed the NMR experiments of Psr_Sp_. TS analysed crystallographic data. PA obtained the secondary structure of Psr_Sp_ based on the TALOS analysis, TS, TA, PA, DL and AA conceptualized the project, supervised different parts of the project.

## COMPETING INTERESTS

The authors declare no competing interests.

## ACKNOWLEDGEMENTS

The authors thank the Swedish NMR Centre for access to the instruments and support. This work was supported by the Swedish Foundation for Strategic Research grant ITM17-0218 to T.A and P.A., grant RSF 23-44-10021 to D.M.L, Swedish Cancer Society (21 1605 Pj 01 H), Cancer och Allergi Fonden (10399), and Swedish Research Council (2021-05061 and 2018-02874 to A.A; 2023-03485 and 2024-06251 to VO).

## DATA AVAILABILITY

Assignment data of Psr_Sp_, with their R1, H2 and NOE relaxations data experimentally obtained and back calculated from free MD simulation together with calculated order parameters S^2^ and population values of conformational ensemble submitted to the BioMagResBank with accession code BMRB ID52556. The structures of trajectory I have been deposited in the Protein Data Bank (PDB) (Entry ID: D_4-G0H2).

## Notes

### Competing Interest Statement

The authors have declared no competing interest.

