## Supplemental information for "Accurate Protein Dynamic Conformational Ensembles: Combining AlphaFold, MD and Amide ^15^N(^1^H) NMR Relaxation"

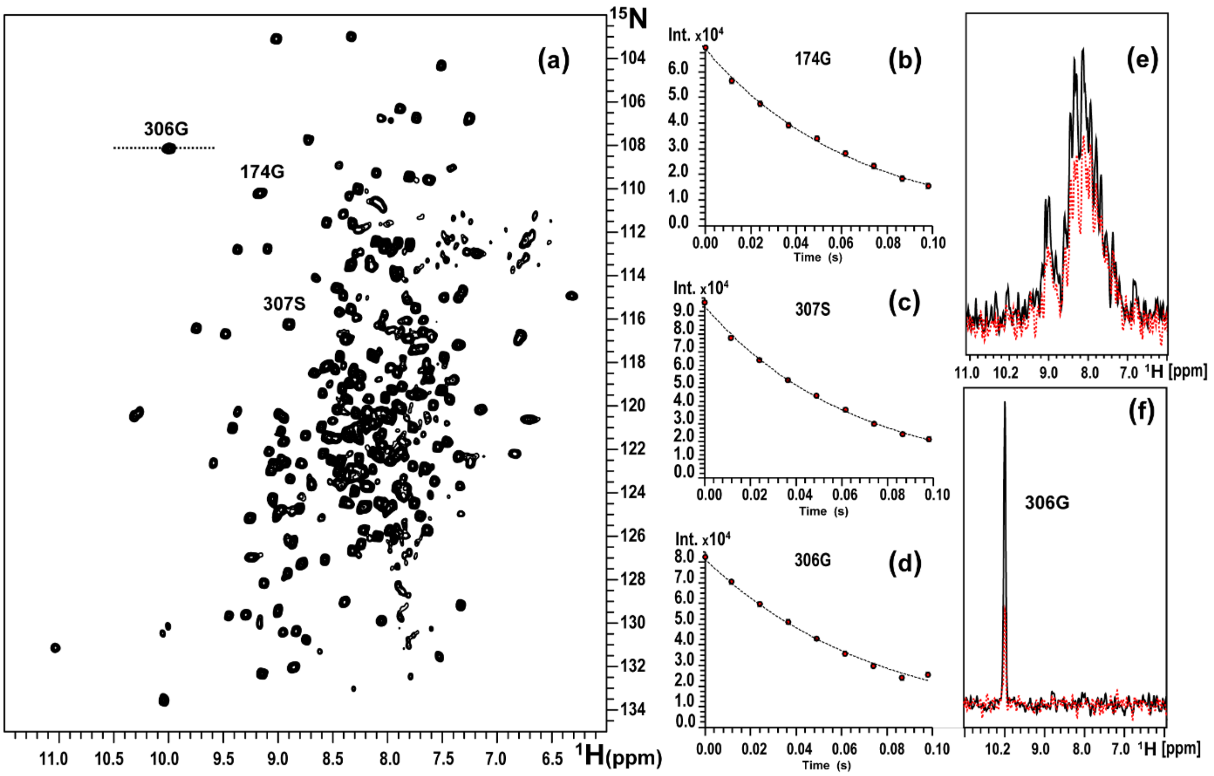


**Figure S1. H2 Spectra of the PsrSp Protein.** (a) First 2D plane at ζ delay equal to 0 of the H2 experiment for the $Psr_{Sp}$ protein; (b), (c), and (d) visualization of intensity changes as a function of ζ delay, fitted with exponential curves to determine the H2 relaxation rates for the selected residues 174G, 307S, and 306G, respectively, as indicated in (a); (e) first increment of the H2 experiment with (black curve) and without (red dashed curve) water flip-back selective pulses, used to prevent saturation of amino proton signals by water; (f) projection through the 306G cross-peak (labeled in (a)), shown as a black curve (first 2D plane) and a red dashed curve (last 2D plane) for ζ delays of 0 and 0.098 s, respectively

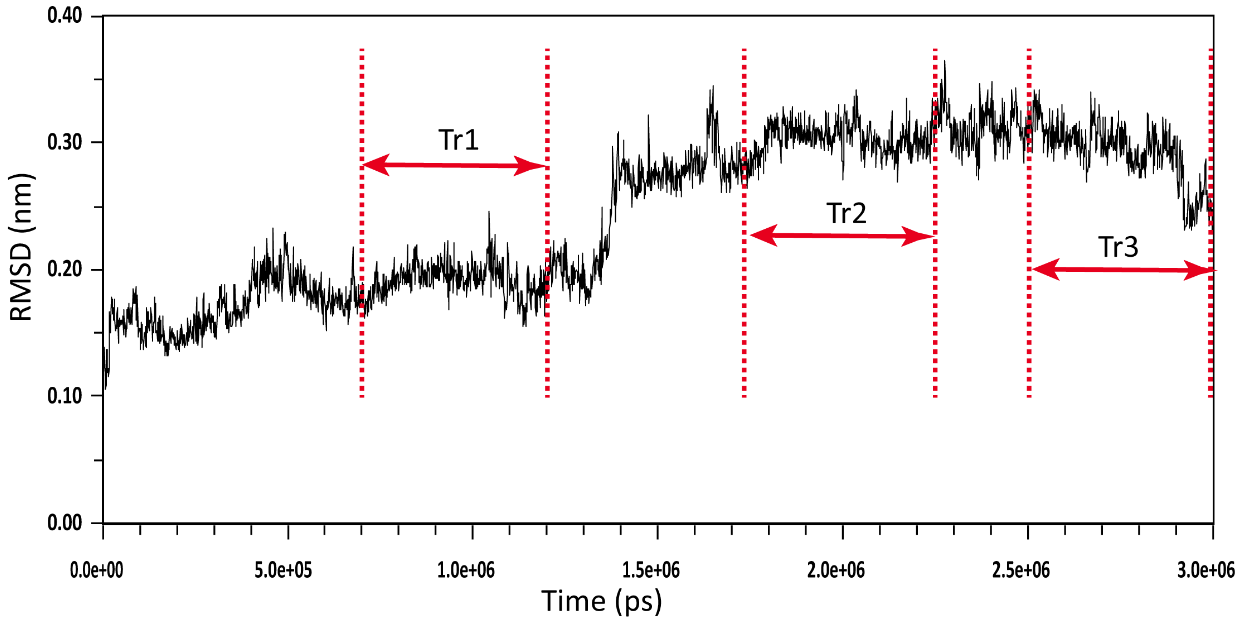


**Figure S2. RMSD along the MD trajectory**.

RMSD vs the initial structure for the backbone heavy atoms of $Psr_{Sp}$ obtained during 3us MD trajectory. The starting structure was obtained from AF of $Psr_{Sp}$. Three trajectory intervals I (700-1200ns), II (1750-2250ns) and III (2500-3000ns) with length 500ns used for back calculations of the relaxation R1, H2, NOE and $S^2$ parameters of protein are depicted by red arrows**.**

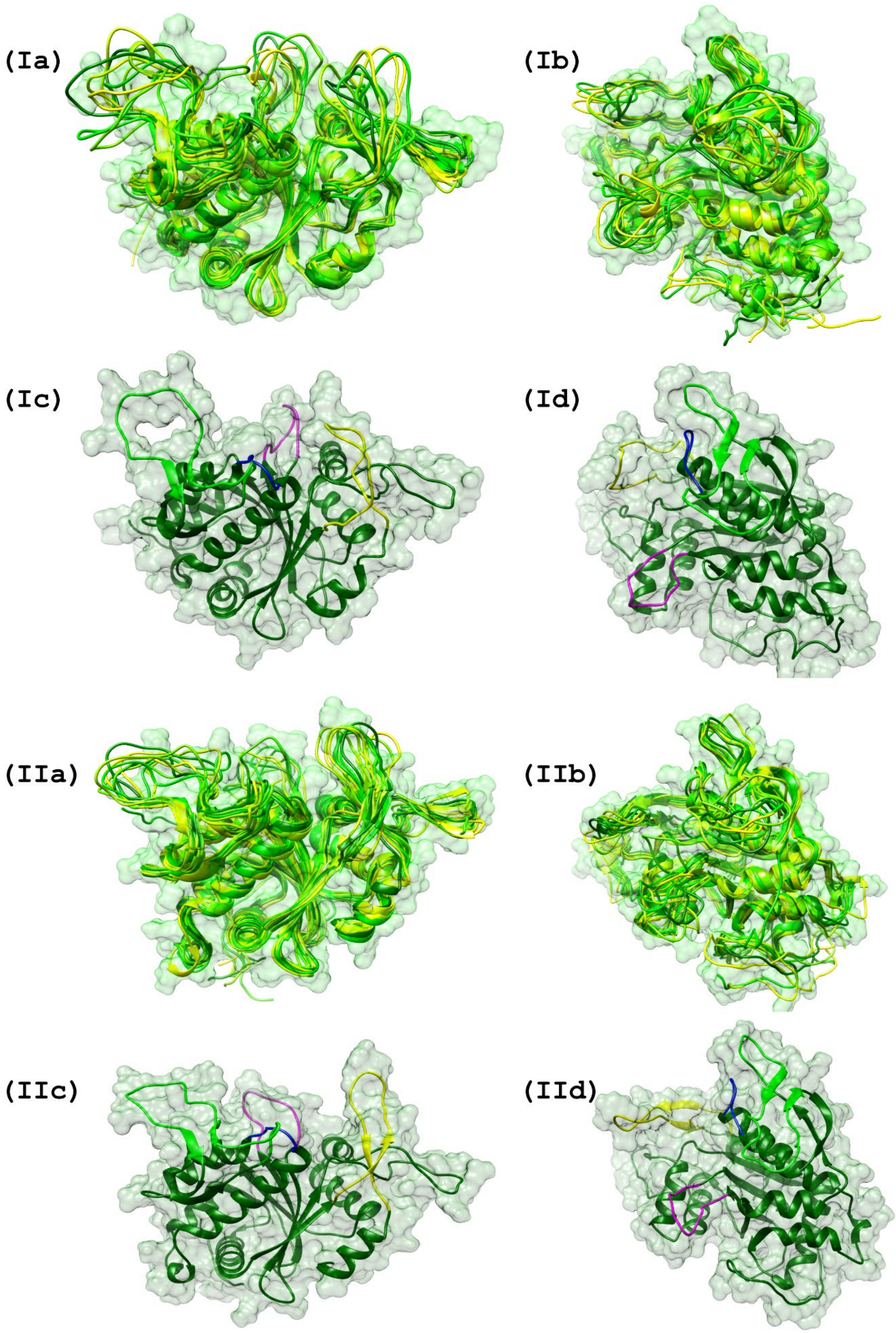
(Ia)
(Ib)
(Ic)
(Id)
(IIa)
(IIb)
(IIc)
(IId)

**Figure S3. Two Conformational Ensembles (I-II) of PsrSp Obtained through 3 μs Long MD Free-Restraint Simulations and Verified by NMR.**

Panels (Ia-d) and (IIa-d) depict ribbon representations of the domain structures of the two ensembles, each comprising 10 models of PsrSp. These models were obtained through clustering analysis of trajectory I (700–1200 ns) and trajectory II (1750–2250 ns), respectively. In panels (**Ia,b**) and (**IIa,b**) the globular $Psr_{Sp}$ domain, spanning residues 130–425, is color-coded according to **Table S2**, with the clusters ranging from highest to lowest populations represented by dark green to yellow, respectively. Panels (**Ic,d**) and (**IIc,d**) illustrate the first structures of the most populated clusters from trajectories I and II, respectively. These structures are displayed in dark green, except for the residues associated with the A, B, C, and D substrate-binding sites of PsrSp, which are highlighted in hot pink, green, dark blue, and yellow, respectively. The structures shown on the right (**Ib,d**) and (**IIb,d**) represent 180-degree rotations of the corresponding structures in panels (**Ia,c**) and (**IIa,c**). The surfaces corresponding to the van der Waals radii of each heteroatom in the PsrSp protein are rendered in transparent white-grey.

**Table S1** Experimentally obtained relaxation parameters R1, H2 and NOE with their errors (σ) of the $Psr_{Sp}$ protein

| N residue | Peak name | R1 ($s^{-1}$) | σ (R1) | NOE | σ (NOE) | H2 ($s^{-1}$) | σ (H2) |
|---|---|---|---|---|---|---|---|
| 130 | 1Gly | 0.837 | 0.036 | 0.690 | 0.078 | 13.92 | 1.80 |
| 131 | 2Glu | | | | | | |
| 132 | 3Val | 1.221 | 0.033 | 0.463 | 0.019 | 8.83 | 0.28 |
| 133 | 4Glu | 1.245 | 0.028 | 0.430 | 0.039 | 8.18 | 0.32 |
| 134 | 5Val | 1.046 | 0.018 | 0.654 | 0.026 | 10.38 | 0.29 |
| 135 | 6Phe | 0.960 | 0.017 | 0.639 | 0.028 | 12.44 | 0.39 |
| 136 | 7Asn | 0.798 | 0.011 | 0.822 | 0.033 | 14.04 | 0.43 |
| 137 | 8Gly | 0.780 | 0.014 | 0.828 | 0.016 | 12.40 | 0.19 |
| 138 | 9Gln | 0.833 | 0.017 | 0.782 | 0.024 | 12.70 | 0.41 |
| 139 | 10Asp | 0.807 | 0.012 | 0.776 | 0.014 | 13.16 | 0.30 |
| 140 | 11Thr | 0.782 | 0.017 | 0.827 | 0.020 | 14.42 | 0.15 |
| 141 | 12Arg | 0.838 | 0.019 | 0.672 | 0.014 | 13.20 | 0.32 |
| 142 | 13Asp | 0.850 | 0.015 | 0.825 | 0.024 | 14.31 | 0.53 |
| 143 | 14Gly | 0.776 | 0.010 | 0.830 | 0.013 | 12.87 | 0.19 |
| 144 | 15Val | 0.791 | 0.020 | 0.840 | 0.023 | 13.99 | 0.29 |
| 145 | 16Asn | 0.830 | 0.024 | 0.819 | 0.021 | 14.09 | 0.39 |
| 146 | 17Ile | 0.799 | 0.028 | 0.873 | 0.034 | 13.45 | 0.52 |
| 147 | 18Leu | 0.815 | 0.025 | 0.857 | 0.038 | 17.52 | 0.53 |
| 148 | 19Ile | 0.764 | 0.041 | 0.792 | 0.059 | 15.40 | 0.90 |
| 149 | 20Met | | | | | | |
| 150 | 21Gly | | | | | | |
| 151 | 22Thr | | | | | | |
| 152 | 23Asp | | | | | | |
| 153 | 24Gly | 1.093 | 0.042 | 0.455 | 0.060 | 8.76 | 0.58 |
| 154 | 25Arg | 1.173 | 0.023 | 0.529 | 0.010 | 8.49 | 0.32 |
| 155 | 26Ile | 1.240 | 0.023 | 0.444 | 0.026 | 7.38 | 0.26 |
| 156 | 27Gly | 1.354 | 0.048 | 0.395 | 0.023 | 6.70 | 0.24 |
| 157 | 28Gln | 1.118 | 0.034 | 0.470 | 0.026 | 7.08 | 0.32 |
| 158 | 29Asn | 1.083 | 0.025 | 0.286 | 0.053 | 4.08 | 0.29 |
| 159 | 30Ser | | | | | | |

| | | | | | | | |
|---|---|---|---|---|---|---|---|
| 160 | 31Val | 0.987 | 0.021 | 0.711 | 0.008 | 11.56 | 0.19 |
| 161 | 32Glu | 1.268 | 0.025 | 0.536 | 0.022 | 8.44 | 0.26 |
| 162 | 33Thr | 1.221 | 0.039 | 0.660 | 0.059 | 10.30 | 1.26 |
| 163 | 34Arg | 0.835 | 0.018 | 0.761 | 0.021 | 13.48 | 0.20 |
| 164 | 35Thr | | | | | | |
| 165 | 36Asp | 0.925 | 0.043 | 0.865 | 0.075 | 14.28 | 0.72 |
| 166 | 37Ser | | | | | | |
| 167 | 38Ile | 0.893 | 0.038 | 0.741 | 0.079 | 19.29 | 3.85 |
| 168 | 39Met | 0.769 | 0.076 | 0.989 | 0.087 | 15.01 | 1.15 |
| 169 | 40Val | 0.799 | 0.022 | 0.821 | 0.052 | 13.07 | 0.85 |
| 170 | 41Leu | 0.919 | 0.047 | 0.985 | 0.090 | 18.57 | 2.11 |
| 171 | 42Asn | | | | | | |
| 172 | 43Val | 0.878 | 0.022 | 0.855 | 0.049 | 12.25 | 1.29 |
| 173 | 44Gly | 0.851 | 0.017 | 0.751 | 0.046 | 12.85 | 0.65 |
| 174 | 45Gly | 0.890 | 0.019 | 0.782 | 0.034 | 14.88 | 0.60 |
| 175 | 46Ser | 0.857 | 0.015 | 0.785 | 0.036 | 13.20 | 0.41 |
| 176 | 47Asp | 0.959 | 0.021 | 0.757 | 0.019 | 14.45 | 0.41 |
| 177 | 48Lys | 0.880 | 0.016 | 0.816 | 0.022 | 15.52 | 0.35 |
| 178 | 49Lys | 0.834 | 0.015 | 0.701 | 0.038 | 13.14 | 0.55 |
| 179 | 50Met | 0.858 | 0.020 | 0.624 | 0.036 | 11.24 | 0.60 |
| 180 | 51Lys | | | | | | |
| 181 | 52Leu | 0.831 | 0.016 | 0.843 | 0.023 | 14.67 | 0.33 |
| 182 | 53Val | | | | | | |
| 183 | 54Ser | | | | | | |
| 184 | 55Phe | 0.884 | 0.029 | 0.668 | 0.033 | 10.08 | 0.45 |
| 185 | 56Met | 0.860 | 0.024 | 0.694 | 0.035 | 9.75 | 0.59 |
| 186 | 57Arg | 0.940 | 0.033 | 0.823 | 0.027 | 14.76 | 0.22 |
| 187 | 58Asp | 0.894 | 0.037 | 0.787 | 0.059 | 14.45 | 1.48 |
| 188 | 59Asn | 0.784 | 0.024 | 0.783 | 0.034 | 15.90 | 0.92 |
| 189 | 60Leu | 0.844 | 0.016 | 0.851 | 0.018 | 14.48 | 0.36 |
| 190 | 61Val | 0.847 | 0.016 | 0.854 | 0.016 | 13.76 | 0.22 |
| 191 | 62Tyr | 0.835 | 0.016 | 0.850 | 0.020 | 14.53 | 0.29 |
| 192 | 63Ile | 0.776 | 0.015 | 0.874 | 0.024 | 16.00 | 0.38 |
| 193 | 64Asp | 0.817 | 0.012 | 0.798 | 0.018 | 14.44 | 0.37 |
| 194 | 65Gly | 0.787 | 0.012 | 0.792 | 0.014 | 12.60 | 0.31 |
| 195 | 66Tyr | 0.828 | 0.017 | 0.817 | 0.023 | 14.00 | 0.72 |
| 196 | 67Ser | 0.685 | 0.016 | 0.617 | 0.026 | 9.33 | 0.25 |
| 197 | 68Gln | 0.782 | 0.010 | 0.708 | 0.016 | 14.40 | 0.27 |
| 198 | 69Val | 0.786 | 0.007 | 0.669 | 0.010 | 12.35 | 0.16 |
| 199 | 70Ile | 0.795 | 0.012 | 0.711 | 0.013 | 13.31 | 0.16 |
| 200 | 71Asn | | | | | | |
| 201 | 72Gly | 0.887 | 0.035 | 0.741 | 0.023 | 11.13 | 0.24 |
| 202 | 73Arg | 0.798 | 0.016 | 0.665 | 0.012 | 14.13 | 0.16 |
| 203 | 74Lys | 0.875 | 0.041 | 0.721 | 0.026 | 13.50 | 0.44 |
| 204 | 75Gln | 0.814 | 0.014 | 0.759 | 0.013 | 13.23 | 0.11 |
| 205 | 76Thr | 0.783 | 0.015 | 0.803 | 0.014 | 12.86 | 0.18 |
| 206 | 77Asp | 0.777 | 0.018 | 0.822 | 0.022 | 13.46 | 0.39 |
| 207 | 78Asn | 0.841 | 0.013 | 0.869 | 0.024 | 14.42 | 0.36 |
| 208 | 79Lys | 0.787 | 0.011 | 0.779 | 0.017 | 12.44 | 0.17 |
| 209 | 80Leu | 0.838 | 0.019 | 0.844 | 0.027 | 13.48 | 0.34 |
| 210 | 81Asn | 1.019 | 0.030 | 0.584 | 0.020 | 10.23 | 0.41 |
| 211 | 82Val | | | | | | |

| | | | | | | | |
|---|---|---|---|---|---|---|---|
| 212 | 83Ala | | | | | | |
| 213 | 84Tyr | | | | | | |
| 214 | 85Glu | 0.949 | 0.041 | 0.730 | 0.052 | 14.74 | 1.50 |
| 215 | 86Leu | 0.908 | 0.022 | 0.826 | 0.026 | 16.32 | 0.47 |
| 216 | 87Gly | 0.874 | 0.019 | 0.863 | 0.020 | 14.80 | 0.48 |
| 217 | 88Glu | 0.898 | 0.016 | 0.816 | 0.033 | 15.30 | 0.69 |
| 218 | 89Gln | 0.907 | 0.014 | 0.723 | 0.015 | 12.74 | 0.25 |
| 219 | 90Glu | 0.884 | 0.015 | 0.745 | 0.017 | 12.84 | 0.28 |
| 220 | 91Gly | 0.906 | 0.016 | 0.206 | 0.016 | 7.82 | 0.15 |
| 221 | 92Gln | 1.025 | 0.021 | 0.569 | 0.022 | 11.54 | 0.36 |
| 222 | 93Lys | 0.897 | 0.018 | 0.735 | 0.021 | 12.53 | 0.34 |
| 223 | 94Gly | 0.822 | 0.065 | 1.107 | 0.124 | 15.52 | 1.54 |
| 224 | 95Ala | 0.937 | 0.025 | 0.743 | 0.048 | 11.03 | 0.66 |
| 225 | 96Glu | | | | | | |
| 226 | 97Met | 0.860 | 0.030 | 0.740 | 0.034 | 15.83 | 0.82 |
| 227 | 98Val | 0.860 | 0.018 | 0.834 | 0.017 | 14.59 | 0.33 |
| 228 | 99Arg | 0.869 | 0.021 | 0.888 | 0.029 | 14.93 | 0.30 |
| 229 | 100Gln | 0.855 | 0.014 | 0.911 | 0.021 | 15.32 | 0.25 |
| 230 | 101Val | 0.811 | 0.009 | 0.901 | 0.021 | 14.87 | 0.42 |
| 231 | 102Leu | 0.890 | 0.025 | 0.843 | 0.023 | 16.66 | 0.42 |
| 232 | 103Lys | 0.906 | 0.023 | 0.801 | 0.030 | 15.82 | 0.42 |
| 233 | 104Asp | 0.839 | 0.015 | 0.826 | 0.024 | 14.86 | 0.75 |
| 234 | 105Asn | | | | | | |
| 235 | 106Phe | | | | | | |
| 236 | 107Asp | 0.886 | 0.023 | 0.851 | 0.047 | 10.83 | 1.42 |
| 237 | 108Leu | 0.841 | 0.022 | 0.829 | 0.019 | 13.40 | 0.34 |
| 238 | 109Asp | 0.854 | 0.023 | 0.801 | 0.025 | 13.38 | 0.36 |
| 239 | 110Ile | 0.816 | 0.016 | 0.818 | 0.027 | 13.49 | 0.45 |
| 240 | 111Lys | 0.859 | 0.018 | 0.825 | 0.026 | 14.59 | 0.67 |
| 241 | 112Tyr | 0.831 | 0.021 | 0.914 | 0.035 | 12.89 | 0.48 |
| 242 | 113Tyr | | | | | | |
| 243 | 114Ala | | | | | | |
| 244 | 115Leu | | | | | | |
| 245 | 116Val | | | | | | |
| 246 | 117Asp | 0.741 | 0.023 | 0.689 | 0.029 | 12.02 | 0.54 |
| 247 | 118Phe | 0.863 | 0.024 | 0.870 | 0.030 | 14.17 | 0.36 |
| 248 | 119Gln | 0.829 | 0.027 | 0.850 | 0.038 | 13.94 | 0.71 |
| 249 | 120Ala | 0.844 | 0.020 | 0.780 | 0.033 | 16.31 | 0.44 |
| 250 | 121Phe | 0.874 | 0.035 | 0.855 | 0.035 | 13.97 | 1.23 |
| 251 | 122Ala | | | | | | |
| 252 | 123Thr | | | | | | |
| 253 | 124Ala | | | | | | |
| 254 | 125 Ile | | | | | | |
| 255 | 126Asp | 0.835 | 0.031 | 0.890 | 0.084 | 13.70 | 2.83 |
| 256 | 127Thr | | | | | | |
| 257 | 128Leu | | | | | | |
| 258 | 129Phe | | | | | | |
| 259 | 130Pro | | | | | | |
| 260 | 131Asp | 0.865 | 0.028 | 0.827 | 0.020 | 12.74 | 0.26 |
| 261 | 132Gly | 0.850 | 0.016 | 0.864 | 0.018 | 14.21 | 0.21 |
| 262 | 133Val | 0.813 | 0.013 | 0.872 | 0.018 | 15.23 | 0.21 |
| 263 | 134Thr | 0.799 | 0.020 | 0.653 | 0.046 | 13.91 | 0.64 |

| | | | | | | | |
|---|---|---|---|---|---|---|---|
| 264 | 135Ile | 0.763 | 0.010 | 0.834 | 0.019 | 16.08 | 0.33 |
| 265 | 136Asp | 0.833 | 0.012 | 0.826 | 0.019 | 14.69 | 0.24 |
| 266 | 137Ala | 0.736 | 0.009 | 0.877 | 0.018 | 15.14 | 0.12 |
| 267 | 138Gln | 0.851 | 0.020 | 0.750 | 0.020 | 13.17 | 0.17 |
| 268 | 139Phe | 0.789 | 0.011 | 0.714 | 0.017 | 11.78 | 0.19 |
| 269 | 140Ser | 0.843 | 0.017 | 0.759 | 0.018 | 12.10 | 0.15 |
| 270 | 141Thr | 0.954 | 0.036 | 0.799 | 0.066 | 8.43 | 1.55 |
| 271 | 142Leu | 0.824 | 0.015 | 0.779 | 0.016 | 14.26 | 0.17 |
| 272 | 143Asn | 0.856 | 0.023 | 0.596 | 0.038 | 14.94 | 0.74 |
| 273 | 144Gly | 0.867 | 0.013 | 0.652 | 0.022 | 12.21 | 0.16 |
| 274 | 145Arg | 0.869 | 0.013 | 0.756 | 0.013 | 14.24 | 0.17 |
| 275 | 146Pro | | | | | | |
| 276 | 147Leu | 0.777 | 0.007 | 0.614 | 0.014 | 12.57 | 0.17 |
| 277 | 148Thr | 0.891 | 0.052 | 0.641 | 0.047 | 11.80 | 1.82 |
| 278 | 149Glu | 0.915 | 0.022 | 0.752 | 0.021 | 15.95 | 0.28 |
| 279 | 150Ala | 0.830 | 0.013 | 0.742 | 0.018 | 14.68 | 0.24 |
| 280 | 151Thr | 0.820 | 0.014 | 0.766 | 0.018 | 13.14 | 0.34 |
| 281 | 152Val | 0.877 | 0.014 | 0.840 | 0.023 | 13.21 | 0.19 |
| 282 | 153Gly | 1.145 | 0.017 | 0.620 | 0.018 | 8.94 | 0.17 |
| 283 | 154Asp | 1.184 | 0.021 | 0.564 | 0.009 | 8.27 | 0.11 |
| 284 | 155Asp | 1.338 | 0.029 | 0.440 | 0.014 | 6.21 | 0.10 |
| 285 | 156Leu | 1.380 | 0.031 | 0.301 | 0.010 | 5.86 | 0.16 |
| 286 | 157Tyr | 1.370 | 0.019 | 0.429 | 0.012 | 6.36 | 0.09 |
| 287 | 158Ala | 1.446 | 0.038 | 0.359 | 0.014 | 5.70 | 0.13 |
| 288 | 159Thr | 1.344 | 0.047 | 0.266 | 0.012 | 4.45 | 0.16 |
| 289 | 160Glu | 1.370 | 0.044 | 0.316 | 0.017 | 4.68 | 0.20 |
| 290 | 161Thr | 1.352 | 0.050 | 0.181 | 0.016 | 3.96 | 0.35 |
| 291 | 162Glu | 1.386 | 0.043 | 0.314 | 0.012 | 4.30 | 0.53 |
| 292 | 163Ser | 1.223 | 0.041 | 0.321 | 0.012 | 5.00 | 0.30 |
| 293 | 164Pro | | | | | | |
| 294 | 165Thr | 1.097 | 0.032 | 0.622 | 0.011 | 7.97 | 0.21 |
| 295 | 166Gln | 1.207 | 0.036 | 0.545 | 0.021 | 9.85 | 0.22 |
| 296 | 167Thr | 0.970 | 0.019 | 0.660 | 0.023 | 12.14 | 0.39 |
| 297 | 168Ile | 0.788 | 0.011 | 0.745 | 0.024 | 13.99 | 0.38 |
| 298 | 169Lys | 0.793 | 0.009 | 0.795 | 0.019 | 14.72 | 0.17 |
| 299 | 170Val | 0.769 | 0.006 | 0.797 | 0.017 | 13.27 | 0.17 |
| 300 | 171Gly | 0.776 | 0.012 | 0.875 | 0.016 | 14.02 | 0.29 |
| 301 | 172Lys | 0.907 | 0.021 | 0.835 | 0.014 | 13.84 | 0.27 |
| 302 | 173Gln | 0.797 | 0.016 | 0.859 | 0.019 | 15.23 | 0.22 |
| 303 | 174Gln | 0.801 | 0.012 | 0.807 | 0.013 | 13.45 | 0.23 |
| 304 | 175Met | 0.778 | 0.010 | 0.844 | 0.021 | 12.52 | 0.26 |
| 305 | 176Asn | 0.833 | 0.014 | 0.788 | 0.014 | 13.68 | 0.12 |
| 306 | 177Gly | 0.795 | 0.011 | 0.852 | 0.031 | 13.34 | 0.17 |
| 307 | 178Ser | 0.807 | 0.012 | 0.851 | 0.030 | 15.55 | 0.39 |
| 308 | 179Thr | 0.903 | 0.019 | 0.640 | 0.027 | 12.18 | 0.52 |
| 309 | 180Leu | 0.862 | 0.017 | 0.857 | 0.031 | 16.52 | 0.34 |
| 310 | 181Leu | 0.813 | 0.015 | 0.822 | 0.037 | 14.94 | 0.54 |
| 311 | 182Asn | 0.808 | 0.017 | 0.840 | 0.047 | 14.47 | 0.92 |
| 312 | 183Tyr | 0.856 | 0.022 | 0.866 | 0.038 | 16.88 | 0.75 |
| 313 | 184Ala | 0.892 | 0.022 | 0.866 | 0.044 | 12.88 | 0.79 |
| 314 | 185Arg | | | | | | |
| 315 | 186Phe | | | | | | |

| | | | | | | | |
|---|---|---|---|---|---|---|---|
| 316 | 187Arg | | | | | | |
| 317 | 188Asp | | | | | | |
| 318 | 189Asp | 0.821 | 0.013 | 0.899 | 0.033 | 14.28 | 0.60 |
| 319 | 190Asp | 0.832 | 0.020 | 0.829 | 0.030 | 12.13 | 0.89 |
| 320 | 191Glu | 0.836 | 0.010 | 0.858 | 0.018 | 16.74 | 0.37 |
| 321 | 192Ala | 0.894 | 0.019 | 0.857 | 0.021 | 16.39 | 0.45 |
| 322 | 193Asp | | | | | | |
| 323 | 194Tyr | | | | | | |
| 324 | 195Gly | 1.025 | 0.056 | 0.786 | 0.083 | 15.30 | 1.88 |
| 325 | 196Arg | 1.180 | 0.047 | 0.366 | 0.035 | 7.41 | 0.65 |
| 326 | 197Thr | | | | | | |
| 327 | 198Lys | | | | | | |
| 328 | 199Arg | 1.323 | 0.031 | 0.095 | 0.036 | 4.57 | 0.17 |
| 329 | 200Gln | 0.917 | 0.017 | 0.666 | 0.019 | 10.99 | 0.29 |
| 330 | 201Gln | | | | | | |
| 331 | 202Gln | 0.853 | 0.029 | 0.800 | 0.051 | 14.64 | 1.58 |
| 332 | 203Val | | | | | | |
| 333 | 204Leu | | | | | | |
| 334 | 205Thr | 0.905 | 0.017 | 0.651 | 0.025 | 12.39 | 0.58 |
| 335 | 206Ala | 0.851 | 0.012 | 0.864 | 0.034 | 15.60 | 0.56 |
| 336 | 207Ile | 0.781 | 0.022 | 0.874 | 0.044 | 16.47 | 0.87 |
| 337 | 208Leu | | | | | | |
| 338 | 209Glu | 0.833 | 0.027 | 0.771 | 0.054 | 15.51 | 0.75 |
| 339 | 210Gln | | | | | | |
| 340 | 211 Ile | 0.850 | 0.017 | 0.822 | 0.049 | 15.94 | 1.45 |
| 341 | 212Lys | 0.930 | 0.031 | 0.782 | 0.047 | 14.09 | 0.92 |
| 342 | 213Asp | | | | | | |
| 343 | 214Pro | | | | | | |
| 344 | 215Thr | 1.063 | 0.027 | 0.000 | 0.000 | 12.01 | 1.31 |
| 345 | 216Lys | | | | | | |
| 346 | 217Leu | | | | | | |
| 347 | 218Phe | | | | | | |
| 348 | 219Thr | | | | | | |
| 349 | 220Gly | | | | | | |
| 350 | 221 Ser | | | | | | |
| 351 | 222Glu | | | | | | |
| 352 | 223Ala | | | | | | |
| 353 | 224Leu | 0.557 | 0.043 | 0.528 | 0.145 | 2.52 | 1.20 |
| 354 | 225Gly | 1.042 | 0.033 | 0.702 | 0.079 | 12.19 | 1.88 |
| 355 | 226Lys | | | | | | |
| 356 | 227Val | | | | | | |
| 357 | 228Phe | | | | | | |
| 358 | 229Ala | | | | | | |
| 359 | 230Met | | | | | | |
| 360 | 231Thr | 1.225 | 0.064 | 0.231 | 0.066 | 2.87 | 0.92 |
| 361 | 232 Ser | 1.113 | 0.047 | 0.724 | 0.080 | 17.02 | 4.18 |
| 362 | 233Thr | 0.803 | 0.018 | 0.839 | 0.042 | 14.60 | 0.98 |
| 363 | 234Asn | | | | | | |
| 364 | 235Val | 0.828 | 0.018 | 0.824 | 0.020 | 14.91 | 0.59 |
| 365 | 236Pro | | | | | | |
| 366 | 237Tyr | 0.831 | 0.016 | 0.825 | 0.036 | 14.89 | 0.45 |
| 367 | 238Thr | 0.924 | 0.074 | 0.717 | 0.119 | 20.88 | 4.20 |

| | | | | | | | |
|---|---|---|---|---|---|---|---|
| 368 | 239Phe | | | | | | |
| 369 | 240Leu | | | | | | |
| 370 | 241Leu | | | | | | |
| 371 | 242Thr | | | | | | |
| 372 | 243Asn | | | | | | |
| 373 | 244Gly | 1.126 | 0.050 | 0.733 | 0.121 | 23.33 | 6.00 |
| 374 | 245Leu | 1.011 | 0.026 | 0.608 | 0.015 | 10.15 | 0.54 |
| 375 | 246 Ser | 0.893 | 0.041 | 0.940 | 0.104 | 12.39 | 1.64 |
| 376 | 247Val | | | | | | |
| 377 | 248Leu | | | | | | |
| 378 | 249Asp | | | | | | |
| 379 | 250Gly | | | | | | |
| 380 | 251Ala | 1.004 | 0.033 | 0.731 | 0.046 | 13.28 | 0.76 |
| 381 | 252Lys | 1.097 | 0.040 | 0.645 | 0.070 | 13.99 | 1.56 |
| 382 | 253Asn | 1.124 | 0.071 | 0.698 | 0.051 | 11.95 | 1.27 |
| 383 | 254Gly | 0.975 | 0.017 | 0.607 | 0.021 | 11.29 | 0.25 |
| 384 | 255 Ile | 1.029 | 0.022 | 0.612 | 0.044 | 9.25 | 0.60 |
| 385 | 256Glu | 0.822 | 0.012 | 0.800 | 0.027 | 13.71 | 0.42 |
| 386 | 257Lys | 0.819 | 0.008 | 0.782 | 0.015 | 12.86 | 0.20 |
| 387 | 258Leu | 0.837 | 0.018 | 0.722 | 0.028 | 12.96 | 0.38 |
| 388 | 259Thr | 0.890 | 0.016 | 0.769 | 0.016 | 14.70 | 0.28 |
| 389 | 260Ile | 0.834 | 0.016 | 0.847 | 0.015 | 12.84 | 0.24 |
| 390 | Pro | | | | | | |
| 391 | 262Glu | 0.823 | 0.019 | 0.818 | 0.028 | 12.73 | 0.53 |
| 392 | 263Leu | 0.776 | 0.031 | 0.726 | 0.053 | 13.67 | 0.45 |
| 393 | 264Gly | 0.791 | 0.039 | 0.565 | 0.063 | 11.07 | 0.58 |
| 394 | 265Asp | 0.884 | 0.017 | 0.728 | 0.016 | 14.59 | 0.37 |
| 395 | 266Trp | 0.775 | 0.012 | 0.834 | 0.020 | 12.92 | 0.37 |
| 396 | 267Val | 0.806 | 0.012 | 0.847 | 0.019 | 13.93 | 0.54 |
| 397 | 268Asp | 0.785 | 0.007 | 0.805 | 0.017 | 13.48 | 0.20 |
| 398 | 269Ala | 0.742 | 0.012 | 0.800 | 0.016 | 14.05 | 0.24 |
| 399 | 270Tyr | 0.753 | 0.008 | 0.667 | 0.011 | 10.57 | 0.22 |
| 400 | 271Asp | 0.774 | 0.014 | 0.793 | 0.016 | 14.73 | 0.18 |
| 401 | 272Val | 1.011 | 0.023 | 0.631 | 0.012 | 10.19 | 0.40 |
| 402 | 273Tyr | 0.960 | 0.018 | 0.700 | 0.010 | 12.28 | 0.53 |
| 403 | 274Gly | 0.817 | 0.010 | 0.792 | 0.019 | 14.10 | 0.31 |
| 404 | 275Gly | 0.847 | 0.013 | 0.826 | 0.020 | 14.54 | 0.23 |
| 405 | 276Leu | 0.844 | 0.023 | 0.685 | 0.011 | 12.51 | 0.20 |
| 406 | 277Gly | 0.758 | 0.012 | 0.729 | 0.017 | 13.14 | 0.26 |
| 407 | 278Leu | 0.822 | 0.015 | 0.882 | 0.016 | 15.08 | 0.23 |
| 408 | 279Leu | 0.797 | 0.011 | 0.857 | 0.017 | 15.81 | 0.29 |
| 409 | 280Val | 0.821 | 0.012 | 0.858 | 0.020 | 13.91 | 0.26 |
| 410 | 281Asp | 0.830 | 0.019 | 0.823 | 0.015 | 14.67 | 0.28 |
| 411 | 282Gln | 0.775 | 0.014 | 0.805 | 0.016 | 14.03 | 0.20 |
| 412 | 283Asn | 0.855 | 0.032 | 0.830 | 0.023 | 15.15 | 0.56 |
| 413 | 284Lys | 0.846 | 0.015 | 0.851 | 0.019 | 15.26 | 0.26 |
| 414 | 285Tyr | 0.846 | 0.016 | 0.859 | 0.022 | 14.87 | 0.18 |
| 415 | 286Gln | 0.819 | 0.014 | 0.816 | 0.017 | 15.36 | 0.36 |
| 416 | 287Thr | 1.050 | 0.025 | 0.611 | 0.010 | 10.14 | 0.29 |
| 417 | 288Lys | 0.826 | 0.017 | 0.854 | 0.015 | 15.69 | 0.40 |
| 418 | 289Leu | 0.818 | 0.017 | 0.865 | 0.018 | 15.30 | 0.34 |
| 419 | 290Ala | 0.818 | 0.013 | 0.864 | 0.012 | 15.69 | 0.29 |

| | | | | | | | |
|---|---|---|---|---|---|---|---|
| 420 | 291Gln | 0.834 | 0.016 | 0.844 | 0.014 | 14.91 | 0.20 |
| 421 | 292Met | 0.793 | 0.010 | 0.837 | 0.021 | 14.16 | 0.12 |
| 422 | 293Gly | 0.793 | 0.016 | 0.809 | 0.020 | 14.06 | 0.27 |
| 423 | 294Leu | 0.888 | 0.017 | 0.863 | 0.024 | 15.40 | 0.58 |
| 424 | 295Arg | 0.851 | 0.016 | 0.864 | 0.017 | 14.33 | 0.26 |

**Table S2** The population of the 10 best clusters obtained from Trajectories I and II by cluster analyse of the $Psr_{Sp}$ protein

| Number of cluster/ type of trajectory | 1 | 2 | 3 | 4 | 5 | 6 | 7 | 8 | 9 | 10 | others |
|---|---|---|---|---|---|---|---|---|---|---|---|
| **Trajectory I[a] 700-1200ns** | 52.9% | 9.3% | 5.8% | 5.0% | 4.6% | 4.3% | 3.8% | 2.3% | 2.0% | 1.4% | 8.6% |
| **Trajectory II[b] 1750-2250ns** | 44.2% | 15.0% | 13.0% | 7.5% | 2.7% | 2.2% | 1.8% | 1.7% | 1.3% | 1.2% | 9.4% |

[a] For trajectory II 700-1200 нс: the cut off cluster analyse was RMSD 0.095nm.
[b] For trajectory II 1750-2250 нс: the cut off cluster analyse was RMSD 0.105nm.

**Table S3** Back calculated relaxation parameters R1, H2, NOE and order parameter $S^2$ with their errors **(σ)** obtained from trajectory **I** (700-1200ns) of the $Psr_{Sp}$ protein

| N residue | Peak name | R1 ($s^{-1}$) | σ (R1) | NOE | σ (NOE) | H2 ($s^{-1}$) | σ (H2) | $S^2$ | σ ($S^2$) |
|---|---|---|---|---|---|---|---|---|---|
| 130 | 1Gly | 0.91 | 0.059 | 0.288 | 0.042 | -8.246 | 0.82 | 0.297 | 0.101 |
| 131 | 2Glu | 0.887 | 0.06 | 0.474 | 0.057 | -9.199 | 0.953 | 0.405 | 0.123 |
| 132 | 3Val | 0.907 | 0.06 | 0.47 | 0.038 | -8.579 | 0.968 | 0.343 | 0.129 |
| 133 | 4Glu | 0.956 | 0.077 | 0.495 | 0.075 | -9.796 | 1.214 | 0.409 | 0.15 |
| 134 | 5Val | 0.918 | 0.11 | 0.661 | 0.039 | -9.445 | 1.022 | 0.438 | 0.114 |
| 135 | 6Phe | 1.004 | 0.091 | 0.675 | 0.03 | -11.614 | 1.183 | 0.629 | 0.109 |
| 136 | 7Asn | 0.892 | 0.038 | 0.678 | 0.015 | -12.06 | 0.712 | 0.699 | 0.035 |
| 137 | 8Gly | 0.795 | 0.043 | 0.346 | 0.074 | -7.273 | 0.46 | 0.403 | 0.003 |
| 138 | 9Gln | 0.883 | 0.033 | 0.767 | 0.009 | -13.228 | 0.763 | 0.814 | 0.015 |
| 139 | 10Asp | 0.853 | 0.032 | 0.783 | 0.007 | -13.278 | 0.759 | 0.831 | 0.006 |
| 140 | 11Thr | 0.818 | 0.031 | 0.767 | 0.013 | -12.622 | 0.731 | 0.795 | 0.008 |
| 141 | 12Arg | 0.871 | 0.032 | 0.744 | 0.01 | -13.27 | 0.758 | 0.831 | 0.004 |
| 142 | 13Asp | 0.919 | 0.036 | 0.74 | 0.01 | -12.357 | 0.732 | 0.641 | 0.061 |
| 143 | 14Gly | 0.864 | 0.032 | 0.759 | 0.021 | -13.034 | 0.777 | 0.804 | 0.019 |
| 144 | 15Val | 0.898 | 0.033 | 0.807 | 0.006 | -14.174 | 0.809 | 0.886 | 0.004 |
| 145 | 16Asn | 0.901 | 0.033 | 0.817 | 0.006 | -14.317 | 0.816 | 0.899 | 0.001 |
| 146 | 17Ile | 0.908 | 0.034 | 0.826 | 0.005 | -14.583 | 0.831 | 0.919 | 5E-04 |
| 147 | 18Leu | 0.909 | 0.034 | 0.829 | 0.006 | -14.587 | 0.832 | 0.92 | 7E-04 |
| 148 | 19Ile | 0.888 | 0.033 | 0.826 | 0.005 | -14.24 | 0.812 | 0.896 | 8E-04 |
| 149 | 20Met | 0.902 | 0.033 | 0.818 | 0.005 | -14.324 | 0.817 | 0.898 | 0.003 |
| 150 | 21Gly | 0.88 | 0.033 | 0.791 | 0.015 | -13.782 | 0.807 | 0.867 | 0.012 |
| 151 | 22Thr | 0.904 | 0.033 | 0.786 | 0.018 | -14.039 | 0.804 | 0.874 | 0.01 |
| 152 | 23Asp | 0.917 | 0.034 | 0.75 | 0.014 | -13.565 | 0.777 | 0.838 | 0.009 |
| 153 | 24Gly | 1.051 | 0.068 | 0.678 | 0.017 | -11.817 | 0.769 | 0.695 | 0.033 |

| | | | | | | | | | |
|---|---|---|---|---|---|---|---|---|---|
| 154 | 25Arg | 0.955 | 0.044 | 0.547 | 0.023 | -9.212 | 0.629 | 0.312 | 0.067 |
| 155 | 26Ile | 1.122 | 0.078 | 0.415 | 0.029 | -7.154 | 0.737 | 0.318 | 0.085 |
| 156 | 27Gly | 1.199 | 0.082 | 0.424 | 0.039 | -6.486 | 0.469 | 0.24 | 0.012 |
| 157 | 28Gln | 1.16 | 0.058 | 0.394 | 0.046 | -5.549 | 0.423 | 0.187 | 0.018 |
| 158 | 29Asn | 1.208 | 0.068 | 0.369 | 0.035 | -5.235 | 0.363 | 0.153 | 0.011 |
| 159 | 30Ser | 1.28 | 0.054 | 0.339 | 0.029 | -4.089 | 0.263 | 0.057 | 0.017 |
| 160 | 31Val | 1.152 | 0.058 | 0.378 | 0.023 | -4.714 | 0.314 | 0.156 | 0.016 |
| 161 | 32Glu | 1.296 | 0.056 | 0.424 | 0.026 | -6.737 | 0.423 | 0.259 | 0.014 |
| 162 | 33Thr | 1.158 | 0.058 | 0.549 | 0.029 | -9.136 | 0.528 | 0.48 | 0.006 |
| 163 | 34Arg | 1.029 | 0.039 | 0.689 | 0.014 | -11.944 | 0.687 | 0.682 | 0.013 |
| 164 | 35Thr | 0.944 | 0.036 | 0.748 | 0.01 | -13.193 | 0.758 | 0.806 | 0.007 |
| 165 | 36Asp | 0.917 | 0.034 | 0.782 | 0.009 | -13.643 | 0.785 | 0.839 | 0.014 |
| 166 | 37Ser | 0.893 | 0.033 | 0.763 | 0.028 | -13.207 | 0.83 | 0.756 | 0.059 |
| 167 | 38Ile | 0.903 | 0.037 | 0.801 | 0.007 | -13.576 | 0.824 | 0.749 | 0.075 |
| 168 | 39Met | 0.914 | 0.034 | 0.826 | 0.005 | -14.6 | 0.832 | 0.921 | 9E-04 |
| 169 | 40 Val | 0.902 | 0.033 | 0.809 | 0.006 | -14.367 | 0.819 | 0.904 | 2E-04 |
| 170 | 41Leu | 0.909 | 0.034 | 0.831 | 0.006 | -14.61 | 0.833 | 0.921 | 4E-04 |
| 171 | 42Asn | 0.893 | 0.033 | 0.823 | 0.005 | -14.303 | 0.815 | 0.901 | 8E-04 |
| 172 | 43Val | 0.903 | 0.033 | 0.822 | 0.006 | -14.364 | 0.819 | 0.902 | 0.002 |
| 173 | 44Gly | 0.863 | 0.032 | 0.818 | 0.005 | -13.579 | 0.774 | 0.855 | 0.002 |
| 174 | 45Gly | 0.86 | 0.032 | 0.799 | 0.006 | -13.44 | 0.767 | 0.844 | 0.003 |
| 175 | 46Ser | 0.91 | 0.035 | 0.749 | 0.019 | -13.456 | 0.783 | 0.835 | 0.012 |
| 176 | 47Asp | 0.823 | 0.031 | 0.732 | 0.014 | -12.198 | 0.699 | 0.761 | 0.006 |
| 177 | 48Lys | 0.887 | 0.033 | 0.728 | 0.013 | -13.021 | 0.751 | 0.809 | 0.008 |
| 178 | 49Lys | 0.867 | 0.032 | 0.697 | 0.017 | -12.47 | 0.74 | 0.778 | 0.012 |
| 179 | 50Met | 0.887 | 0.033 | 0.776 | 0.008 | -13.542 | 0.773 | 0.841 | 0.005 |
| 180 | 51Lys | 0.877 | 0.033 | 0.792 | 0.008 | -13.594 | 0.775 | 0.853 | 8E-04 |
| 181 | 52Leu | 0.907 | 0.034 | 0.821 | 0.005 | -14.444 | 0.823 | 0.91 | 8E-04 |
| 182 | 53Val | 0.905 | 0.033 | 0.824 | 0.005 | -14.41 | 0.821 | 0.907 | 4E-04 |
| 183 | 54Ser | 0.903 | 0.033 | 0.82 | 0.006 | -14.346 | 0.818 | 0.903 | 0.001 |
| 184 | 55Phe | 0.905 | 0.033 | 0.826 | 0.006 | -14.435 | 0.823 | 0.908 | 0.002 |
| 185 | 56Met | 0.908 | 0.034 | 0.806 | 0.007 | -14.038 | 0.801 | 0.875 | 0.004 |
| 186 | 57Arg | 0.902 | 0.033 | 0.807 | 0.006 | -13.977 | 0.797 | 0.871 | 0.003 |
| 187 | 58Asp | 0.912 | 0.034 | 0.823 | 0.005 | -14.425 | 0.822 | 0.908 | 0.001 |
| 188 | 59Asn | 0.905 | 0.033 | 0.819 | 0.005 | -14.309 | 0.816 | 0.9 | 7E-04 |
| 189 | 60Leu | 0.903 | 0.033 | 0.817 | 0.006 | -14.228 | 0.811 | 0.894 | 0.003 |
| 190 | 61Val | 0.902 | 0.033 | 0.807 | 0.007 | -14.19 | 0.809 | 0.892 | 5E-04 |
| 191 | 62Tyr | 0.908 | 0.034 | 0.798 | 0.007 | -13.935 | 0.796 | 0.872 | 0.003 |
| 192 | 63Ile | 0.911 | 0.034 | 0.782 | 0.01 | -13.988 | 0.801 | 0.879 | 0.005 |
| 193 | 64Asp | 0.899 | 0.033 | 0.778 | 0.009 | -13.762 | 0.785 | 0.863 | 0.001 |
| 194 | 65Gly | 0.809 | 0.03 | 0.751 | 0.01 | -12.192 | 0.695 | 0.761 | 0.002 |
| 195 | 66Tyr | 0.81 | 0.042 | 0.724 | 0.017 | -9.776 | 0.56 | 0.572 | 0.007 |
| 196 | 67Ser | 0.892 | 0.038 | 0.54 | 0.036 | -10.506 | 0.657 | 0.615 | 0.018 |
| 197 | 68Gln | 0.867 | 0.033 | 0.706 | 0.01 | -12.071 | 0.721 | 0.747 | 0.014 |
| 198 | 69Val | 0.918 | 0.035 | 0.64 | 0.018 | -11.122 | 0.741 | 0.673 | 0.025 |
| 199 | 70Ile | 0.994 | 0.067 | 0.636 | 0.021 | -11.832 | 0.808 | 0.718 | 0.036 |
| 200 | 71Asn | 0.903 | 0.034 | 0.611 | 0.029 | -11.805 | 0.687 | 0.728 | 0.01 |
| 201 | 72Gly | 0.893 | 0.034 | 0.624 | 0.026 | -11.483 | 0.665 | 0.704 | 0.008 |
| 202 | 73Arg | 0.889 | 0.04 | 0.595 | 0.015 | -10.933 | 0.652 | 0.668 | 0.014 |
| 203 | 74Lys | 0.916 | 0.034 | 0.602 | 0.015 | -11.885 | 0.68 | 0.734 | 0.003 |
| 204 | 75Gln | 0.894 | 0.033 | 0.619 | 0.018 | -11.746 | 0.704 | 0.723 | 0.015 |
| 205 | 76Thr | 0.909 | 0.046 | 0.563 | 0.021 | -10.768 | 0.663 | 0.61 | 0.026 |

| | | | | | | | | | |
|---|---|---|---|---|---|---|---|---|---|
| 206 | 77Asp | 0.887 | 0.033 | 0.606 | 0.074 | -12.082 | 0.852 | 0.735 | 0.033 |
| 207 | 78Asn | 0.905 | 0.034 | 0.755 | 0.019 | -12.987 | 0.801 | 0.792 | 0.023 |
| 208 | 79Lys | 0.895 | 0.033 | 0.808 | 0.006 | -13.819 | 0.788 | 0.867 | 0.002 |
| 209 | 80Leu | 0.913 | 0.034 | 0.821 | 0.006 | -14.371 | 0.819 | 0.905 | 4E-04 |
| 210 | 81Asn | 0.915 | 0.034 | 0.821 | 0.006 | -14.485 | 0.826 | 0.909 | 8E-04 |
| 211 | 82Val | 0.91 | 0.034 | 0.823 | 0.006 | -14.397 | 0.821 | 0.904 | 0.002 |
| 212 | 83Ala | 0.911 | 0.034 | 0.828 | 0.005 | -14.532 | 0.828 | 0.915 | 6E-04 |
| 213 | 84Tyr | 0.917 | 0.034 | 0.826 | 0.006 | -14.491 | 0.826 | 0.908 | 0.001 |
| 214 | 85Glu | 0.927 | 0.034 | 0.821 | 0.006 | -14.469 | 0.825 | 0.906 | 0.001 |
| 215 | 86Leu | 0.929 | 0.034 | 0.822 | 0.006 | -14.621 | 0.833 | 0.919 | 4E-04 |
| 216 | 87Gly | 0.911 | 0.034 | 0.821 | 0.006 | -14.34 | 0.817 | 0.896 | 0.001 |
| 217 | 88Glu | 0.922 | 0.034 | 0.81 | 0.007 | -14.374 | 0.82 | 0.899 | 0.002 |
| 218 | 89Gln | 0.921 | 0.035 | 0.713 | 0.012 | -13.576 | 0.776 | 0.846 | 0.002 |
| 219 | 90Glu | 0.896 | 0.034 | 0.725 | 0.012 | -13.234 | 0.758 | 0.821 | 0.003 |
| 220 | 91Gly | 0.856 | 0.035 | 0.531 | 0.032 | -10.012 | 0.655 | 0.598 | 0.02 |
| 221 | 92Gln | 0.994 | 0.041 | 0.518 | 0.026 | -10.889 | 0.65 | 0.625 | 0.02 |
| 222 | 93Lys | 0.94 | 0.036 | 0.656 | 0.014 | -12.116 | 0.697 | 0.728 | 0.008 |
| 223 | 94Gly | 0.91 | 0.034 | 0.771 | 0.007 | -13.962 | 0.796 | 0.871 | 0.002 |
| 224 | 95Ala | 0.895 | 0.033 | 0.799 | 0.006 | -14.023 | 0.799 | 0.878 | 0.002 |
| 225 | 96Glu | 0.912 | 0.034 | 0.822 | 0.006 | -14.5 | 0.827 | 0.911 | 0.002 |
| 226 | 97Met | 0.925 | 0.034 | 0.832 | 0.005 | -14.784 | 0.843 | 0.932 | 6E-04 |
| 227 | 98Val | 0.919 | 0.034 | 0.829 | 0.005 | -14.701 | 0.838 | 0.925 | 8E-04 |
| 228 | 99Arg | 0.925 | 0.034 | 0.832 | 0.005 | -14.809 | 0.844 | 0.932 | 0.002 |
| 229 | 100Gln | 0.923 | 0.034 | 0.83 | 0.005 | -14.816 | 0.845 | 0.933 | 8E-04 |
| 230 | 101Val | 0.922 | 0.034 | 0.832 | 0.005 | -14.784 | 0.843 | 0.932 | 5E-04 |
| 231 | 102Leu | 0.926 | 0.034 | 0.835 | 0.005 | -14.817 | 0.845 | 0.934 | 4E-04 |
| 232 | 103Lys | 0.922 | 0.034 | 0.832 | 0.005 | -14.705 | 0.838 | 0.927 | 9E-04 |
| 233 | 104Asp | 0.919 | 0.034 | 0.831 | 0.005 | -14.646 | 0.835 | 0.924 | 4E-04 |
| 234 | 105Asn | 0.892 | 0.033 | 0.807 | 0.007 | -13.906 | 0.793 | 0.875 | 0.002 |
| 235 | 106Phe | 0.88 | 0.033 | 0.81 | 0.006 | -13.785 | 0.786 | 0.868 | 9E-04 |
| 236 | 107Asp | 0.892 | 0.033 | 0.813 | 0.006 | -14.058 | 0.802 | 0.884 | 0.002 |
| 237 | 108Leu | 0.825 | 0.031 | 0.721 | 0.013 | -12.256 | 0.7 | 0.768 | 0.002 |
| 238 | 109Asp | 0.89 | 0.033 | 0.808 | 0.008 | -14.104 | 0.805 | 0.887 | 0.002 |
| 239 | 110Ile | 0.882 | 0.033 | 0.803 | 0.009 | -13.72 | 0.789 | 0.845 | 0.014 |
| 240 | 111Lys | 0.894 | 0.033 | 0.817 | 0.006 | -14.06 | 0.802 | 0.875 | 0.004 |
| 241 | 112Tyr | 0.882 | 0.033 | 0.824 | 0.006 | -13.967 | 0.799 | 0.878 | 0.005 |
| 242 | 113Tyr | 0.912 | 0.034 | 0.829 | 0.005 | -14.615 | 0.833 | 0.919 | 0.002 |
| 243 | 114Ala | 0.909 | 0.034 | 0.83 | 0.006 | -14.559 | 0.83 | 0.918 | 0.001 |
| 244 | 115Leu | 0.895 | 0.033 | 0.826 | 0.006 | -14.276 | 0.814 | 0.892 | 0.005 |
| 245 | 116Val | 0.908 | 0.034 | 0.819 | 0.008 | -14.312 | 0.817 | 0.888 | 0.008 |
| 246 | 117Asp | 0.894 | 0.033 | 0.804 | 0.007 | -13.886 | 0.793 | 0.864 | 0.003 |
| 247 | 118Phe | 0.902 | 0.033 | 0.816 | 0.006 | -14.339 | 0.817 | 0.901 | 7E-04 |
| 248 | 119Gln | 0.894 | 0.033 | 0.79 | 0.007 | -14.01 | 0.799 | 0.874 | 0.002 |
| 249 | 120Ala | 0.861 | 0.032 | 0.798 | 0.007 | -13.544 | 0.772 | 0.846 | 0.002 |
| 250 | 121Phe | 0.866 | 0.032 | 0.799 | 0.009 | -13.529 | 0.774 | 0.847 | 0.003 |
| 251 | 122Ala | 0.916 | 0.034 | 0.829 | 0.006 | -14.684 | 0.837 | 0.923 | 8E-04 |
| 252 | 123Thr | 0.907 | 0.033 | 0.819 | 0.006 | -14.487 | 0.826 | 0.912 | 4E-04 |
| 253 | 124Ala | 0.909 | 0.034 | 0.825 | 0.006 | -14.565 | 0.83 | 0.917 | 7E-04 |
| 254 | 125 Ile | 0.921 | 0.034 | 0.828 | 0.005 | -14.739 | 0.84 | 0.927 | 4E-04 |
| 255 | 126Asp | 0.917 | 0.034 | 0.828 | 0.005 | -14.661 | 0.836 | 0.922 | 3E-04 |
| 256 | 127Thr | 0.874 | 0.032 | 0.819 | 0.006 | -13.982 | 0.798 | 0.881 | 0.002 |
| 257 | 128Leu | 0.908 | 0.034 | 0.821 | 0.005 | -14.418 | 0.822 | 0.903 | 0.002 |

| | | | | | | | | | |
|---|---|---|---|---|---|---|---|---|---|
| 258 | 129Phe | 0.907 | 0.034 | 0.821 | 0.006 | -14.354 | 0.818 | 0.9 | 3E-04 |
| 259 | 130Pro | | | | | | | | |
| 260 | 131Asp | 0.859 | 0.032 | 0.799 | 0.008 | -13.142 | 0.75 | 0.821 | 9E-04 |
| 261 | 132Gly | 0.839 | 0.031 | 0.808 | 0.006 | -13.058 | 0.745 | 0.819 | 9E-04 |
| 262 | 133Val | 0.911 | 0.034 | 0.789 | 0.008 | -13.473 | 0.774 | 0.843 | 0.006 |
| 263 | 134Thr | 0.923 | 0.035 | 0.81 | 0.007 | -13.617 | 0.778 | 0.852 | 0.002 |
| 264 | 135Ile | 0.914 | 0.034 | 0.817 | 0.007 | -14.103 | 0.806 | 0.884 | 0.004 |
| 265 | 136Asp | 0.894 | 0.033 | 0.787 | 0.008 | -13.392 | 0.769 | 0.832 | 0.008 |
| 266 | 137Ala | 0.927 | 0.038 | 0.757 | 0.014 | -12.686 | 0.724 | 0.774 | 0.003 |
| 267 | 138Gln | 0.922 | 0.034 | 0.757 | 0.011 | -13.3 | 0.762 | 0.825 | 0.003 |
| 268 | 139Phe | 0.909 | 0.034 | 0.722 | 0.012 | -12.582 | 0.72 | 0.776 | 0.004 |
| 269 | 140Ser | 0.949 | 0.036 | 0.766 | 0.011 | -13.484 | 0.77 | 0.828 | 0.004 |
| 270 | 141Thr | 0.931 | 0.065 | 0.698 | 0.022 | -10.28 | 1.207 | 0.591 | 0.071 |
| 271 | 142Leu | 0.991 | 0.059 | 0.777 | 0.008 | -13.521 | 0.848 | 0.836 | 0.026 |
| 272 | 143Asn | 0.933 | 0.037 | 0.678 | 0.014 | -12.826 | 0.732 | 0.792 | 0.002 |
| 273 | 144Gly | 0.936 | 0.038 | 0.727 | 0.012 | -13.211 | 0.757 | 0.805 | 0.002 |
| 274 | 145Arg | 0.853 | 0.038 | 0.764 | 0.008 | -12.005 | 0.692 | 0.74 | 0.005 |
| 275 | 146Pro | | | | | | | | |
| 276 | 147Leu | 0.936 | 0.046 | 0.717 | 0.017 | -10.091 | 0.731 | 0.379 | 0.078 |
| 277 | 148Thr | 0.927 | 0.039 | 0.771 | 0.011 | -12.495 | 0.723 | 0.68 | 0.03 |
| 278 | 149Glu | 0.878 | 0.033 | 0.754 | 0.008 | -13.07 | 0.747 | 0.8 | 0.003 |
| 279 | 150Ala | 0.904 | 0.034 | 0.781 | 0.009 | -13.538 | 0.778 | 0.85 | 0.006 |
| 280 | 151Thr | 0.862 | 0.035 | 0.745 | 0.016 | -12.403 | 0.726 | 0.773 | 0.013 |
| 281 | 152Val | 0.962 | 0.054 | 0.722 | 0.024 | -12.673 | 0.845 | 0.76 | 0.039 |
| 282 | 153Gly | 1.056 | 0.046 | 0.61 | 0.086 | -9.727 | 0.632 | 0.282 | 0.035 |
| 283 | 154Asp | 1.121 | 0.053 | 0.437 | 0.059 | -7.88 | 0.584 | 0.377 | 0.021 |
| 284 | 155Asp | 1.306 | 0.059 | 0.242 | 0.039 | -5.148 | 0.402 | 0.096 | 0.012 |
| 285 | 156Leu | 1.365 | 0.058 | 0.244 | 0.055 | -5.348 | 0.422 | 0.122 | 0.021 |
| 286 | 157Tyr | 1.366 | 0.056 | 0.221 | 0.032 | -4.609 | 0.372 | 0.038 | 0.013 |
| 287 | 158Ala | 1.209 | 0.054 | 0.139 | 0.06 | -4.696 | 0.383 | 0.059 | 0.033 |
| 288 | 159Thr | 1.18 | 0.049 | 0.148 | 0.071 | -3.584 | 0.264 | 0.043 | 0.005 |
| 289 | 160Glu | 1.284 | 0.059 | 0.124 | 0.057 | -3.913 | 0.27 | 0.072 | 0.018 |
| 290 | 161Thr | 1.097 | 0.049 | 0.088 | 0.054 | -4.377 | 0.346 | 0.026 | 0.011 |
| 291 | 162Glu | 1.14 | 0.064 | 0.055 | 0.034 | -4.51 | 0.433 | 0.031 | 0.013 |
| 292 | 163Ser | 1.081 | 0.06 | 0.14 | 0.037 | -5.557 | 0.548 | 0.165 | 0.021 |
| 293 | 164Pro | | | | | | | | |
| 294 | 165Thr | 1.141 | 0.085 | 0.332 | 0.085 | -7.431 | 0.924 | 0.217 | 0.029 |
| 295 | 166Gln | 1.275 | 0.143 | 0.52 | 0.047 | -8.795 | 1.512 | 0.446 | 0.127 |
| 296 | 167Thr | 1.129 | 0.092 | 0.672 | 0.019 | -11.115 | 0.996 | 0.66 | 0.059 |
| 297 | 168Ile | 0.925 | 0.038 | 0.735 | 0.013 | -13.112 | 0.757 | 0.81 | 0.007 |
| 298 | 169Lys | 0.893 | 0.038 | 0.707 | 0.01 | -12.558 | 0.718 | 0.766 | 0.004 |
| 299 | 170Val | 0.924 | 0.038 | 0.76 | 0.016 | -13.202 | 0.789 | 0.811 | 0.019 |
| 300 | 171Gly | 0.909 | 0.035 | 0.75 | 0.011 | -12.806 | 0.735 | 0.784 | 0.004 |
| 301 | 172Lys | 0.879 | 0.033 | 0.74 | 0.013 | -12.747 | 0.729 | 0.782 | 0.002 |
| 302 | 173Gln | 0.907 | 0.034 | 0.749 | 0.019 | -13.306 | 0.798 | 0.826 | 0.017 |
| 303 | 174Gln | 0.888 | 0.033 | 0.767 | 0.014 | -13.025 | 0.787 | 0.794 | 0.017 |
| 304 | 175Met | 0.887 | 0.033 | 0.78 | 0.007 | -12.983 | 0.74 | 0.809 | 9E-04 |
| 305 | 176Asn | 0.888 | 0.033 | 0.818 | 0.006 | -14.054 | 0.802 | 0.886 | 0.002 |
| 306 | 177Gly | 0.914 | 0.034 | 0.825 | 0.005 | -14.628 | 0.834 | 0.921 | 5E-04 |
| 307 | 178Ser | 0.91 | 0.034 | 0.826 | 0.006 | -14.565 | 0.83 | 0.917 | 0.001 |
| 308 | 179Thr | 0.903 | 0.033 | 0.821 | 0.006 | -14.438 | 0.823 | 0.907 | 3E-04 |
| 309 | 180Leu | 0.926 | 0.034 | 0.832 | 0.005 | -14.876 | 0.848 | 0.937 | 3E-04 |

| | | | | | | | | | |
|---|---|---|---|---|---|---|---|---|---|
| 310 | 181Leu | 0.921 | 0.034 | 0.83 | 0.005 | -14.772 | 0.842 | 0.93 | 4E-04 |
| 311 | 182Asn | 0.921 | 0.034 | 0.825 | 0.005 | -14.755 | 0.841 | 0.929 | 5E-04 |
| 312 | 183Tyr | 0.923 | 0.034 | 0.833 | 0.005 | -14.83 | 0.845 | 0.934 | 2E-04 |
| 313 | 184Ala | 0.92 | 0.034 | 0.834 | 0.005 | -14.761 | 0.841 | 0.929 | 4E-04 |
| 314 | 185Arg | 0.903 | 0.033 | 0.828 | 0.005 | -14.436 | 0.823 | 0.908 | 0.001 |
| 315 | 186Phe | 0.877 | 0.032 | 0.808 | 0.008 | -13.702 | 0.786 | 0.85 | 0.001 |
| 316 | 187Arg | 0.909 | 0.034 | 0.805 | 0.006 | -14.194 | 0.809 | 0.889 | 5E-04 |
| 317 | 188Asp | 0.886 | 0.033 | 0.806 | 0.007 | -13.717 | 0.783 | 0.856 | 0.003 |
| 318 | 189Asp | 0.851 | 0.033 | 0.172 | 0.098 | -9.237 | 0.588 | 0.552 | 0.014 |
| 319 | 190Asp | 0.935 | 0.035 | 0.761 | 0.011 | -14.032 | 0.8 | 0.88 | 0.002 |
| 320 | 191Glu | 0.915 | 0.037 | 0.729 | 0.011 | -12.85 | 0.734 | 0.782 | 0.003 |
| 321 | 192Ala | 0.91 | 0.034 | 0.796 | 0.007 | -14.053 | 0.802 | 0.883 | 0.003 |
| 322 | 193Asp | 0.894 | 0.033 | 0.821 | 0.005 | -14.051 | 0.801 | 0.883 | 0.001 |
| 323 | 194Tyr | 0.915 | 0.034 | 0.828 | 0.005 | -14.536 | 0.829 | 0.914 | 0.001 |
| 324 | 195Gly | 0.916 | 0.034 | 0.826 | 0.005 | -14.552 | 0.829 | 0.915 | 0.002 |
| 325 | 196Arg | 0.924 | 0.034 | 0.83 | 0.005 | -14.686 | 0.837 | 0.922 | 0.001 |
| 326 | 197Thr | 0.919 | 0.034 | 0.829 | 0.005 | -14.676 | 0.837 | 0.923 | 8E-04 |
| 327 | 198Lys | 0.922 | 0.034 | 0.83 | 0.005 | -14.748 | 0.841 | 0.929 | 5E-04 |
| 328 | 199Arg | 0.916 | 0.034 | 0.831 | 0.005 | -14.659 | 0.836 | 0.923 | 4E-04 |
| 329 | 200Gln | 0.917 | 0.034 | 0.818 | 0.008 | -14.516 | 0.828 | 0.913 | 3E-04 |
| 330 | 201Gln | 0.915 | 0.034 | 0.832 | 0.005 | -14.668 | 0.836 | 0.924 | 1E-04 |
| 331 | 202Gln | 0.912 | 0.034 | 0.828 | 0.005 | -14.589 | 0.832 | 0.919 | 2E-04 |
| 332 | 203Val | 0.923 | 0.034 | 0.83 | 0.005 | -14.771 | 0.842 | 0.93 | 3E-04 |
| 333 | 204Leu | 0.924 | 0.034 | 0.834 | 0.005 | -14.779 | 0.842 | 0.932 | 3E-04 |
| 334 | 205Thr | 0.916 | 0.034 | 0.828 | 0.006 | -14.668 | 0.836 | 0.923 | 5E-04 |
| 335 | 206Ala | 0.92 | 0.034 | 0.828 | 0.005 | -14.724 | 0.839 | 0.927 | 7E-04 |
| 336 | 207Ile | 0.915 | 0.034 | 0.831 | 0.005 | -14.642 | 0.835 | 0.923 | 5E-04 |
| 337 | 208Leu | 0.913 | 0.034 | 0.828 | 0.005 | -14.586 | 0.832 | 0.918 | 0.001 |
| 338 | 209Glu | 0.909 | 0.034 | 0.819 | 0.006 | -14.393 | 0.82 | 0.903 | 6E-04 |
| 339 | 210Gln | 0.906 | 0.034 | 0.811 | 0.008 | -14.011 | 0.799 | 0.875 | 0.001 |
| 340 | 211 Ile | 0.912 | 0.034 | 0.788 | 0.009 | -13.439 | 0.766 | 0.828 | 0.002 |
| 341 | 212Lys | 0.895 | 0.033 | 0.713 | 0.018 | -12.331 | 0.722 | 0.75 | 0.013 |
| 342 | 213Asp | 0.899 | 0.04 | 0.734 | 0.012 | -12.053 | 0.74 | 0.735 | 0.026 |
| 343 | 214Pro | | | | | | | | |
| 344 | 215Thr | 0.914 | 0.034 | 0.79 | 0.007 | -13.797 | 0.787 | 0.857 | 0.003 |
| 345 | 216Lys | 0.909 | 0.034 | 0.762 | 0.009 | -13.198 | 0.758 | 0.813 | 0.009 |
| 346 | 217Leu | 0.924 | 0.035 | 0.783 | 0.011 | -13.521 | 0.773 | 0.826 | 0.007 |
| 347 | 218Phe | 0.947 | 0.035 | 0.797 | 0.008 | -13.908 | 0.794 | 0.839 | 0.009 |
| 348 | 219Thr | 0.944 | 0.036 | 0.747 | 0.014 | -12.57 | 0.737 | 0.759 | 0.012 |
| 349 | 220Gly | 1.198 | 0.066 | 0.537 | 0.045 | -8.43 | 0.935 | 0.294 | 0.081 |
| 350 | 221 Ser | 1.134 | 0.092 | 0.386 | 0.08 | -7.122 | 0.439 | 0.215 | 0.023 |
| 351 | 222Glu | 0.945 | 0.037 | 0.699 | 0.016 | -12.598 | 0.726 | 0.765 | 0.005 |
| 352 | 223Ala | 0.955 | 0.039 | 0.71 | 0.021 | -12.9 | 0.78 | 0.795 | 0.019 |
| 353 | 224Leu | 0.925 | 0.034 | 0.795 | 0.007 | -14.054 | 0.805 | 0.855 | 0.004 |
| 354 | 225Gly | 0.913 | 0.034 | 0.805 | 0.009 | -14.064 | 0.803 | 0.876 | 0.003 |
| 355 | 226Lys | 0.909 | 0.034 | 0.819 | 0.006 | -14.151 | 0.809 | 0.879 | 0.003 |
| 356 | 227Val | 0.924 | 0.035 | 0.81 | 0.007 | -14.184 | 0.809 | 0.873 | 0.004 |
| 357 | 228Phe | 0.928 | 0.035 | 0.814 | 0.007 | -14.354 | 0.819 | 0.887 | 0.003 |
| 358 | 229Ala | 0.893 | 0.034 | 0.801 | 0.008 | -13.71 | 0.783 | 0.848 | 0.003 |
| 359 | 230Met | 0.902 | 0.036 | 0.741 | 0.012 | -13.223 | 0.755 | 0.785 | 0.009 |
| 360 | 231Thr | 0.872 | 0.033 | 0.782 | 0.009 | -13.337 | 0.762 | 0.834 | 0.002 |
| 361 | 232 Ser | 0.873 | 0.032 | 0.787 | 0.009 | -13.425 | 0.766 | 0.837 | 0.003 |

| | | | | | | | | | |
|---|---|---|---|---|---|---|---|---|---|
| 362 | 233Thr | 0.885 | 0.033 | 0.808 | 0.011 | -13.981 | 0.8 | 0.864 | 0.012 |
| 363 | 234Asn | 0.892 | 0.033 | 0.823 | 0.007 | -14.119 | 0.806 | 0.872 | 0.011 |
| 364 | 235Val | 0.903 | 0.033 | 0.828 | 0.005 | -14.301 | 0.816 | 0.892 | 0.004 |
| 365 | 236Pro | | | | | | | | |
| 366 | 237Tyr | 0.928 | 0.035 | 0.809 | 0.01 | -13.708 | 0.794 | 0.776 | 0.037 |
| 367 | 238Thr | 0.954 | 0.036 | 0.815 | 0.007 | -13.675 | 0.79 | 0.713 | 0.043 |
| 368 | 239Phe | 0.909 | 0.034 | 0.817 | 0.006 | -14.206 | 0.81 | 0.886 | 0.001 |
| 369 | 240Leu | 0.934 | 0.035 | 0.815 | 0.006 | -14.131 | 0.806 | 0.843 | 0.008 |
| 370 | 241Leu | 0.932 | 0.035 | 0.822 | 0.005 | -14.227 | 0.812 | 0.854 | 0.008 |
| 371 | 242Thr | 0.919 | 0.034 | 0.782 | 0.009 | -13.558 | 0.784 | 0.815 | 0.013 |
| 372 | 243Asn | 0.938 | 0.038 | 0.786 | 0.012 | -12.865 | 0.766 | 0.703 | 0.033 |
| 373 | 244Gly | 0.882 | 0.033 | 0.809 | 0.006 | -13.517 | 0.771 | 0.823 | 0.005 |
| 374 | 245Leu | 0.913 | 0.034 | 0.806 | 0.006 | -14.034 | 0.801 | 0.866 | 0.003 |
| 375 | 246 Ser | 0.923 | 0.035 | 0.813 | 0.006 | -13.998 | 0.801 | 0.854 | 0.009 |
| 376 | 247Val | 0.92 | 0.034 | 0.821 | 0.005 | -14.501 | 0.827 | 0.905 | 0.003 |
| 377 | 248Leu | 0.92 | 0.034 | 0.814 | 0.006 | -14.46 | 0.824 | 0.904 | 0.003 |
| 378 | 249Asp | 0.925 | 0.034 | 0.814 | 0.006 | -14.483 | 0.826 | 0.904 | 0.002 |
| 379 | 250Gly | 0.91 | 0.034 | 0.819 | 0.007 | -14.414 | 0.822 | 0.902 | 0.002 |
| 380 | 251Ala | 0.916 | 0.034 | 0.818 | 0.007 | -14.51 | 0.827 | 0.911 | 5E-04 |
| 381 | 252Lys | 0.914 | 0.034 | 0.816 | 0.006 | -14.437 | 0.823 | 0.907 | 0.001 |
| 382 | 253Asn | 0.886 | 0.033 | 0.817 | 0.006 | -13.947 | 0.796 | 0.874 | 0.002 |
| 383 | 254Gly | 0.88 | 0.033 | 0.769 | 0.01 | -13.033 | 0.751 | 0.802 | 0.011 |
| 384 | 255 Ile | 0.896 | 0.034 | 0.776 | 0.011 | -13.183 | 0.762 | 0.813 | 0.015 |
| 385 | 256Glu | 0.905 | 0.033 | 0.791 | 0.007 | -14.166 | 0.808 | 0.891 | 9E-04 |
| 386 | 257Lys | 0.863 | 0.032 | 0.806 | 0.006 | -13.612 | 0.776 | 0.855 | 0.001 |
| 387 | 258Leu | 0.858 | 0.032 | 0.801 | 0.006 | -13.419 | 0.765 | 0.84 | 0.001 |
| 388 | 259Thr | 0.875 | 0.032 | 0.813 | 0.006 | -13.772 | 0.786 | 0.868 | 0.002 |
| 389 | 260Ile | 0.912 | 0.034 | 0.823 | 0.005 | -14.473 | 0.825 | 0.911 | 0.001 |
| 390 | Pro | | | | | | | | |
| 391 | 262Glu | 0.945 | 0.035 | 0.752 | 0.01 | -13.081 | 0.747 | 0.803 | 0.002 |
| 392 | 263Leu | 0.948 | 0.035 | 0.708 | 0.011 | -12.774 | 0.729 | 0.779 | 0.002 |
| 393 | 264Gly | 0.877 | 0.033 | 0.674 | 0.015 | -12.228 | 0.709 | 0.753 | 0.008 |
| 394 | 265Asp | 0.909 | 0.034 | 0.774 | 0.008 | -13.64 | 0.778 | 0.852 | 7E-04 |
| 395 | 266Trp | 0.925 | 0.036 | 0.7 | 0.025 | -11.917 | 0.686 | 0.731 | 0.003 |
| 396 | 267Val | 0.941 | 0.035 | 0.733 | 0.01 | -13.264 | 0.757 | 0.819 | 0.002 |
| 397 | 268Asp | 0.964 | 0.043 | 0.655 | 0.016 | -11.193 | 0.649 | 0.64 | 0.011 |
| 398 | 269Ala | 0.959 | 0.068 | 0.498 | 0.033 | -7.755 | 0.573 | 0.198 | 0.06 |
| 399 | 270Tyr | 0.928 | 0.064 | 0.461 | 0.077 | -7.638 | 0.511 | 0.302 | 0.049 |
| 400 | 271Asp | 1.047 | 0.091 | 0.419 | 0.034 | -8.261 | 0.555 | 0.387 | 0.034 |
| 401 | 272Val | 1.148 | 0.094 | 0.318 | 0.05 | -7.126 | 0.505 | 0.3 | 0.035 |
| 402 | 273Tyr | 1.147 | 0.086 | 0.349 | 0.065 | -7.28 | 0.611 | 0.287 | 0.061 |
| 403 | 274Gly | 1.136 | 0.103 | 0.295 | 0.054 | -5.87 | 0.791 | 0.188 | 0.042 |
| 404 | 275Gly | 1.129 | 0.101 | 0.408 | 0.051 | -5.886 | 0.502 | 0.101 | 0.018 |
| 405 | 276Leu | 1.055 | 0.069 | 0.475 | 0.021 | -8.772 | 0.597 | 0.312 | 0.026 |
| 406 | 277Gly | 1.1 | 0.108 | 0.532 | 0.028 | -8.38 | 0.76 | 0.248 | 0.037 |
| 407 | 278Leu | 0.897 | 0.034 | 0.753 | 0.008 | -13.02 | 0.744 | 0.8 | 0.004 |
| 408 | 279Leu | 0.895 | 0.033 | 0.771 | 0.008 | -13.452 | 0.769 | 0.839 | 0.004 |
| 409 | 280Val | 0.923 | 0.035 | 0.747 | 0.01 | -13.164 | 0.76 | 0.814 | 0.004 |
| 410 | 281Asp | 0.947 | 0.035 | 0.701 | 0.011 | -12.77 | 0.73 | 0.788 | 0.001 |
| 411 | 282Gln | 0.931 | 0.035 | 0.669 | 0.01 | -12.306 | 0.702 | 0.756 | 0.003 |
| 412 | 283Asn | 0.951 | 0.039 | 0.665 | 0.015 | -11.839 | 0.682 | 0.697 | 0.017 |
| 413 | 284Lys | 0.942 | 0.036 | 0.698 | 0.017 | -12.343 | 0.705 | 0.764 | 0.003 |

| | | | | | | | | | |
|---|---|---|---|---|---|---|---|---|---|
| 414 | 285Tyr | 0.92 | 0.034 | 0.776 | 0.01 | -13.874 | 0.792 | 0.871 | 0.003 |
| 415 | 286Gln | 0.907 | 0.033 | 0.818 | 0.006 | -14.198 | 0.809 | 0.893 | 5E-04 |
| 416 | 287Thr | 0.917 | 0.034 | 0.813 | 0.006 | -14.267 | 0.813 | 0.896 | 4E-04 |
| 417 | 288Lys | 0.913 | 0.034 | 0.796 | 0.006 | -14.076 | 0.802 | 0.882 | 0.001 |
| 418 | 289Leu | 0.927 | 0.034 | 0.818 | 0.006 | -14.533 | 0.829 | 0.913 | 4E-04 |
| 419 | 290Ala | 0.924 | 0.034 | 0.816 | 0.006 | -14.47 | 0.825 | 0.91 | 7E-04 |
| 420 | 291Gln | 0.919 | 0.034 | 0.809 | 0.006 | -14.343 | 0.818 | 0.901 | 6E-04 |
| 421 | 292Met | 0.903 | 0.033 | 0.803 | 0.006 | -14.032 | 0.8 | 0.881 | 5E-04 |
| 422 | 293Gly | 0.881 | 0.033 | 0.734 | 0.01 | -12.714 | 0.725 | 0.782 | 0.002 |
| 423 | 294Leu | 0.911 | 0.035 | 0.726 | 0.012 | -12.862 | 0.741 | 0.79 | 0.006 |
| 424 | 295Arg | 0.896 | 0.036 | 0.688 | 0.012 | -11.652 | 0.717 | 0.575 | 0.07 |
